# Multimodal substrate recruitment enables CTLH^MKLN1^ E3 ligase to target N-, C-, and internal degrons

**DOI:** 10.64898/2026.09.10.750665

**Authors:** Sara Sepic, Karthik V. Gottemukkala, Jiale Du, Samuel A. Maiwald, Jakub Chrustowicz, Eleftheria C. Papadopoulou, Susanne von Gronau, Brenda A. Schulman

## Abstract

The GID/CTLH family of E3 ubiquitin ligases employs several substrate receptor subunits that recruit distinct degrons, but the substrate recognition mechanism of the CTLH^MKLN1^ assembly has largely remained elusive. Here, we reconstitute CTLH^MKLN1^-dependent ubiquitylation of three biochemically distinct substrates - MKLN1 itself, ZMYND19, and FAM72A-recruited UNG2, and determine cryo-EM structures of each substrate bound to MKLN1. The structures reveal how MKLN1’s discoidin and Kelch β-propeller domains engage substrates through multivalent contacts. MKLN1 recruits itself through discoidin - Kelch interactions between MKLN1 dimers, forming an assembly that competes with other substrates. ZMYND19 and FAM72A each bind to MKLN1 through loops engaging the discoidin trench domain while their C-terminal Arg residues engage the MKLN1 Kelch central channel, identifying MKLN1 as an Arg/C-degron receptor. The acetylated N-terminus of UNG2 is positioned within a Y-shaped tunnel of FAM72A, defining an adaptor-mediated Ac/N-degron recognition mechanism. Thus, our data reveal how combinatorial deployment of MKLN1 discoidin and Kelch domains enable a single receptor subunit to recognize a diverse substrate repertoire, adding to a complex picture of degron recognition across the GID/CTLH family.

## Introduction

Specificity in the ubiquitin system depends on E3 ligases selecting substrates and positioning them for targeting by the E3–E2∼ubiquitin catalytic assembly. The prevailing view is that specificity is achieved by an E3’s dedicated substrate-binding site docking with a cognate degron conserved across its substrates^1,2^. Notably, a common mode of protein targeting by the ubiquitin system involves E3 ligase recognition of N- or C-terminal sequences, termed N- or C-degrons, respectively ^3–14^. For homodimeric E3s or E3 complexes, a pair of substrate-binding domains can engage a pair of homologous degron motifs from a single ubiquitylation target, an oligomer, or two individual substrate molecules^15–21^. Although this mechanistic framework has been useful, it increasingly appears incomplete. An emerging concept from biochemical studies is that some E3s harbor multiple distinct domains or regions that combinatorially allow various modes of engagement to enable ubiquitylation of diverse substrates^22–25^. Understanding how such E3 ligase complexes recognize and orient diverse substrates for ubiquitin targeting remains an important problem in ubiquitin biology.

A particularly sophisticated mode for substrate engagement is utilized by the family of E3 ligases in the GID/CTLH family (yeast/higher eukaryotic nomenclature). GID/CTLH E3s are modular, multiprotein complexes built around a core assembly^26–32^.The GID/CTLH core contains a heterodimeric RING-like catalytic module and a multiprotein scaffold^26,30,33,34^. Initial studies in yeast showed that exchangeable, metabolically-induced substrate receptor subunits bind the core GID complex (via an adaptor protein) and recruit various N-degron sequences in metabolic enzymes^5,29,30,35–41^. An additional subunit, Gid7, forms a dimer that in turn promote dimerization of the core assembly into a giant oval “Chelator” E3 complex^15^. In this giant oval configuration, the E3 complex contains two opposing copies of the GID E3 elements: two catalytic modules, two scaffolds, two Gid7 dimer-mediated assembly interfaces and two substrate receptors bound to adaptors. Structural studies revealed that this oval architecture enables substrate engagement in a manner analogous to an organometallic chelator binding a ligand through multiple points of contacts. The two opposing Gid4 substrate receptor subunits simultaneously capture Pro/N-degrons from two protomers of the tetrameric substrate, Fbp1. This arrangement drives ubiquitylation of specific Fbp1 lysines that are adjacent to metabolite-binding sites^15^. It was initially thought that the homologous mammalian CTLH E3s would function similarly^31,42,43^ by relying on the GID4 substrate receptor to recruit N-degrons^44–46^. However, subsequent work demonstrated that some CTLH complexes lack the GID4 subunit and instead rely on higher eukaryotic Gid7 paralogs to recruit substrates^42,47,48^. Indeed, the human CTLH subunit WDR26, which is homologous to yeast Gid7 and likewise forms a dimer that mediates formation of a giant oval E3 complex, was proposed to be a GID4-independent substrate receptor^15,48–50^.

Unexpectedly, considering that all known yeast GID E3s recruit N-degrons, WDR26’s WD40 domain was found to recognize internal basic degron motifs in the metabolic enzyme substrate NMNAT1^48^. The pair of propellers from adjacent WDR26 subunits can also bind YPEL5. which inhibits substrates from binding to WDR26. YPEL5 has a CRBN-like Yippee domain that can also mediate targeted protein degradation^51^. Thus, mammalian CTLH complexes can deploy both GID4 and WDR26 for substrate recruitment, with the pair of WDR26 subunits also binding a single Yippee domain protein at their substrate-binding sites^45,46,48,49,51–53^.

A different higher eukaryotic subunit, Muskelin (MKLN1), also occupies a Gid7-like architectural position within an oval CTLH E3 assembly^54^. MKLN1 was proposed as yet another substrate-binding receptor^47,55^. However, MKLN1 markedly differs in domain composition, oligomerization properties, and functional output compared to yeast Gid7 and human WDR26. MKLN1 lacks WD40 repeats, and instead uniquely displays discoidin and Kelch domains^15,56–59^, suggesting it may employ entirely distinct mechanisms of substrate engagement. Consistent with this notion, MKLN1-dependent CTLH complexes have been implicated in a wide range of biological processes beyond the turnover of metabolic enzymes. Deletion of MKLN1 in mice leads to deficiencies ranging from behavioral changes and synaptic defects to impaired somatic hypermutation and class switch recombination^54,60,61^. MKLN1 mutations in dogs were found to be the cause of lethal acrodermatitis, a disease characterized by severe skin lesions, immunodeficiency, and growth failure^62^. In cellular systems, MKLN1 depletion perturbs cytoskeletal organization and adhesion^63–65^, while studies in flies suggested roles in developmental transitions^47,55^. This breadth of phenotypes implies that MKLN1-dependent CTLH complexes act on a diverse set of substrates. Yet, despite reports of many proteins becoming stabilized upon mutation of MKLN1, and high-resolution cryo-EM data for MKLN1 itself, structural details of CTLH substrate recruitment dependent on MKLN1 have not been experimentally elucidated.

One reason for this gap in our understanding is that MKLN1-dependent ubiquitylation has only been reconstituted with entirely purified reagents for a single substrate, the DNA repair enzyme UNG2, which is recruited to the FAM72A adaptor to regulate antibody diversification^54,66^. A low resolution cryo-EM map of human CTLH^MKLN1^-FAM72A-UNG2 showed MKLN can organize a large oval CTLH E3 assembly similar to that observed with yeast Gid7 and human WDR26^15,48,54^. However, the available structural data do not permit independent placement of CTLH^MKLN1^ E3 subunits, nor do they reveal details of substrate recognition via the FAM72A adaptor. Although two preprints posted on bioRXiv during preparation of our manuscript suggested MKLN1 recognizes Arg/C-degrons, those studies lack experimental structural data and do not explain how such substrate recruitment places targets relative to the CTLH E3 assembly or the ubiquitylation active sites^67,68^. Furthermore, the existing studies do not explain how MKLN1 itself is subject to ubiquitylation^46,69,70^. As a result, our understanding of the molecular principles that govern substrate recruitment by MKLN1-containing E3 ligases remains limited. Thus, to define structural mechanisms and relative roles of the unique MKLN1 domains in the regulation of distinct substrates, we obtained cryo-EM data for stable proxies of ubiquitylation intermediates, in the form of substrate-bound CTLH E3 complexes with an E2∼ubiquitin conjugate. The cryo-EM maps, taken together with biochemical and cell-based data, reveal a diversity of mechanisms for CTLH^MKLN1^ placement of various substrates for ubiquitylation.

## Results

### Biochemical reconstitution of diverse MKLN1-dependent substrates

As a first step towards defining molecular principles underlying substrate recognition by CTLH^MKLN1^, we used purified reagents to biochemically reconstitute MKLN1-dependent ubiquitylation activity toward previously defined substrates with distinct properties. Of all the subunits of the CTLH complex (Fig. 1a), MKLN1 is the one that is most frequently reported as stabilized upon elimination of CTLH E3 catalytic activity^46,69–72^. ZMYND19 was selected because the E3 subunit directly mediating its recognition is subject to debate^49,69,71^. Meanwhile, UNG2 degradation was shown to depend on the MKLN1-binding adaptor protein FAM72A^54,61^.

**Fig. 1.**
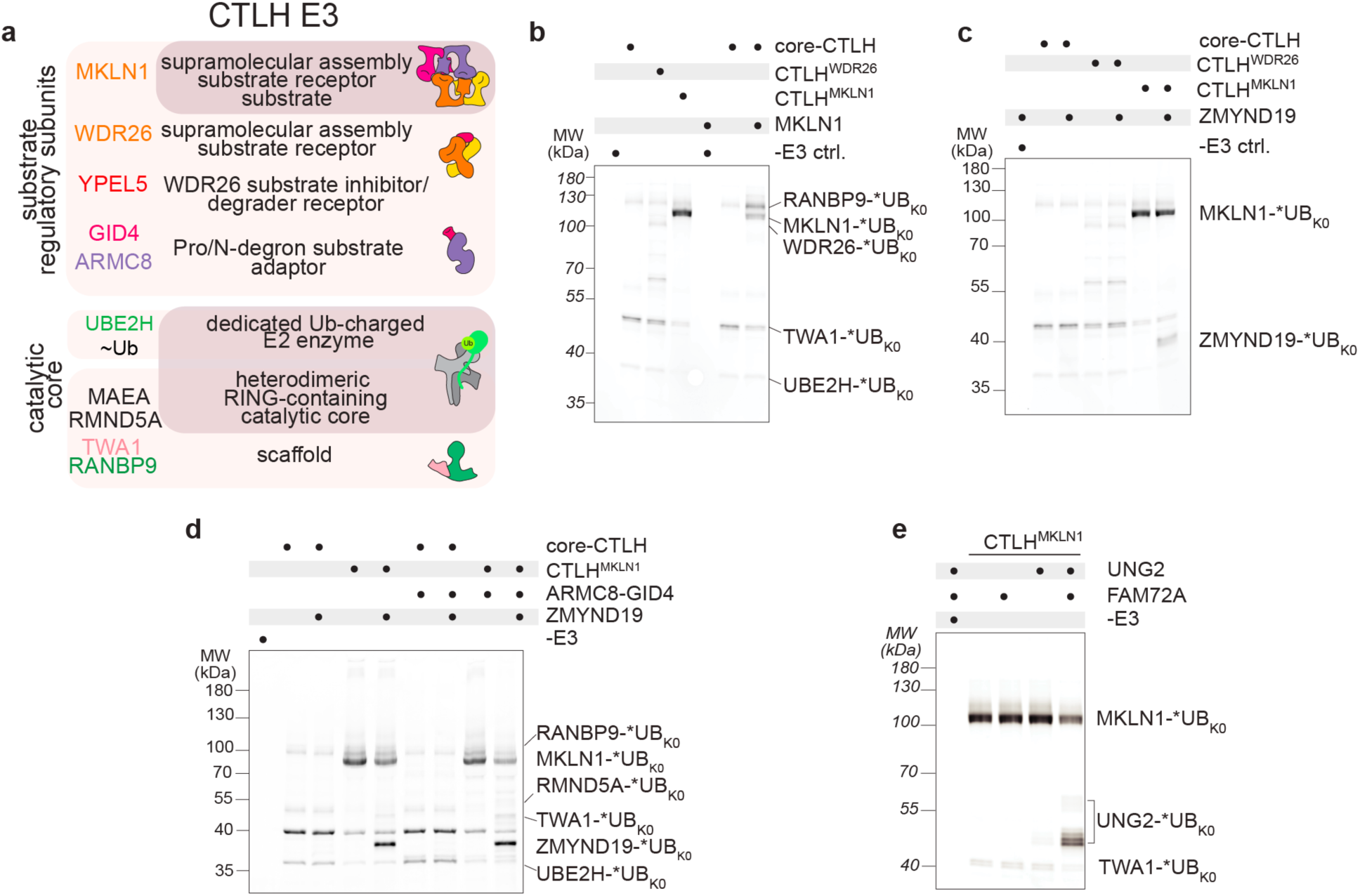
Biochemical reconstitution of CTLH^MKLN1^-dependent substrate ubiquitylation. **a** Schematic of the CTLH E3 complex subunits organized by functional roles. **b** Assays testing MKLN1 ubiquitylation by the indicated CTLH E3 complexes. Ubiquitylation assays used N-terminally fluorescently labeled K0-ubiquitin (*UB_K0_) with all lysines mutated to arginines to visualize direct substrate modification without chain formation, with UBE2H as the E2. **c** In vitro ubiquitylation assays testing ZMYND19 ubiquitylation by the indicated CTLH E3 complexes. **d** Assays testing effect of the GID4 substrate receptor on CTLH^MKLN1^-dependent ZMYND19 ubiquitylation. **e** Assays testing UNG2 ubiquitylation by CTLH^MKLN1^ in the presence or absence of substrate adaptor FAM72A. b–e, All assays were performed for 15 min (n = 2 technically independent experiments).

Experiments compared activity with a “core” CTLH complex (containing RANBP9, TWA1, MAEA, and RMND5A) in the absence or presence of either human Gid7-like subunit WDR26 or MKLN1. We used N-terminally fluorescently-labeled “K0” ubiquitin with all lysines mutated to arginines to visualize direct substrate modification without additional banding from chain formation. Ubiquitylation was achieved by the CTLH E3’s cognate E2, UBE2H^31,73^. MKLN1 was robustly ubiquitylated in our assays, regardless of whether it was co-expressed with the core CTLH E3 complex or added separately (Fig. 1b). Notably, this differs from the other Gid7 ortholog, WDR26, which showed little ubiquitylation. Our assays showed ZMYND19 was ubiquitylated by CTLH^MKLN1^, but not by core-CTLH or CTLH^WDR^^26^ complexes. ZMYND19 ubiquitylation was also unaffected by the GID4 substrate receptor (Fig. 1c, d). Thus, our biochemical data differ from a previous claim that GID4 recruits ZMYND19, but are consistent with prior cellular studies suggesting that MKLN1 is both a ubiquitylation substrate and a substrate receptor for ZMYND19^49,69^. Meanwhile, our assays recapitulate prior work showing that UNG2 ubiquitylation required CTLH^MKLN1^ and the adaptor protein FAM72A^54^ (Fig. 1e).

### Substrate recognition by CTLH^MKLN1^

To determine the structural basis for substrate recognition by the CTLH^MKLN1^ E3, we obtained cryo-EM data for complexes representing ubiquitylation of the three substrates, MKLN1, ZMYND19, and FAM72A-UNG2. The substrate-bound E3 complexes included a stable mimic of the UBE2H∼ubiquitin conjugate. Ubiquitin transfer was prevented by use of a non-reactive complex where ubiquitin’s C-terminus was isopeptide-bonded to a Lys replacement for the UBE2H catalytic Cys^73^. Low resolution cryo-EM maps showed all three substrates in the center of the oval CTLH^MKLN1^ E3, proximal to the UBE2H active sites (Fig. 2a). Details of substrate interactions with the MKLN1 discoidin and Kelch domains, described below, were obtained by fitting the map with structures, either achieved by focused refinements of the complete complex, or from acquiring data for subcomplexes (see Methods, Supplementary Data Table 1, Supplementary Fig. 5-10).

**Fig. 2.**
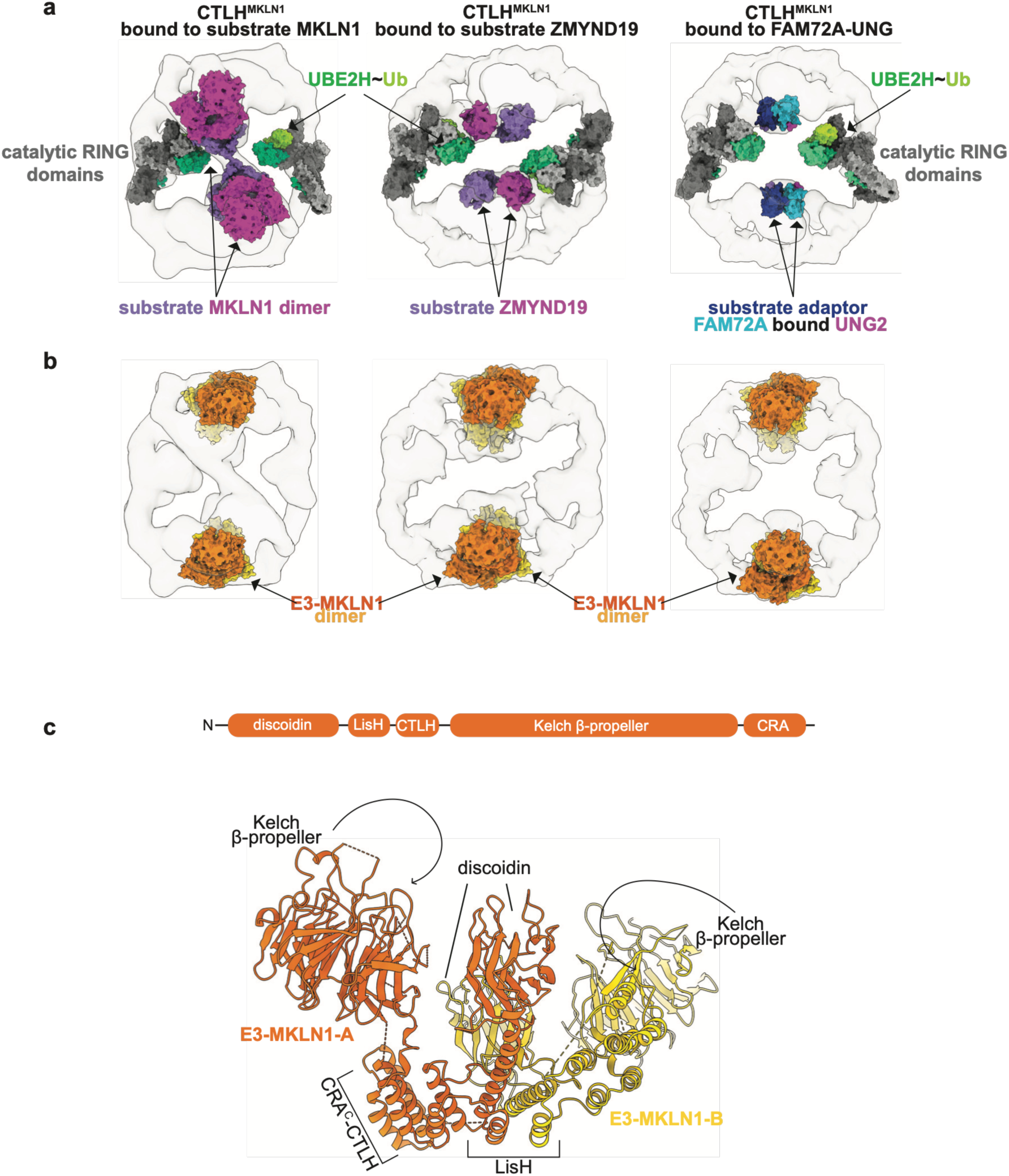
MKLN1 recruits diverse substrates to the center of an oval CTLH assembly. **a** Low-resolution cryo-EM maps of CTLH^MKLN1^-UBE2H∼ubiquitin conjugate bound to substrate MKLN1 (left), substrate ZMYND19 (middle), and FAM72A–UNG2 (right), with fitted structures from this study and the catalytic module (PDB 8PJN) displayed as surface representations. The UBE2H∼ ubiquitin conjugate represents a stable mimic of the ubiquitylation intermediate, in which ubiquitin’s C-terminus is isopeptide-bonded to a Lys replacement for the UBE2H catalytic Cys, preventing ubiquitin transfer. Substrates and substrate adaptor are indicated. **b** Cryo-EM maps with fitted atomic models as surface representations, highlighting the E3-MKLN1 dimer (orange and yellow) within each of the three complexes shown in **a** illustrating the conserved oval assembly architecture. **c** Top: domain architecture of MKLN1. Bottom: structure of E3-MKLN1-A (dark orange) and E3-MKLN1-B (yellow) protomers (this study) fitted into the CTLH^MKLN1^ E3-MKLN1 dimer density, depicting the overall E3-MKLN1 dimer architecture, with domains labelled.

The CTLH^MKLN1^ E3s resemble Chelator-GID and CTLH^WDR^^26^ E3 complexes ^15,48^, in that a pair of Gid7 orthologs (here MKLN1) homodimerize to close the oval-shaped E3 ligase assembly (Fig. 2b, Supplementary Fig. 1a). Gid7 and WDR26 form oval E3s through their LisH, CTLH, and WD40 beta-propeller domains, where the LisH domain mediates homodimerization, and the CTLH domain (together with other domains) mediates interactions with core E3 subunits^30,58,59,74^. MKLN1 is characterized by LisH, CTLH, discoidin, and Kelch β-propeller domains (Fig. 2c). The former two domains function analogously to those in Gid7 and WDR26, while the latter two domains mediate substrate-binding in various ways described below.

### Substrate: MKLN1

The complex with MKLN1 differed strikingly from prior structures of Chelator-GID or CTLH^WDR^^26^ E3 complexes, which feature a hollow cavity for binding to substrates in the center of their oval shape (Supplementary Fig. 1b). Here, however, the oval was filled by two additional MKLN1 protomers (Fig. 3a). The structural data suggest how MKLN1 recruits itself as a substrate: while one MKLN1 dimer (hereafter “E3-MKLN1”) integrates into the CTLH oval assembly, the second MKLN1 dimer (hereafter “Sub-MKLN1”) is positioned as a substrate (Supplementary Fig. 2a).

**Fig. 3.**
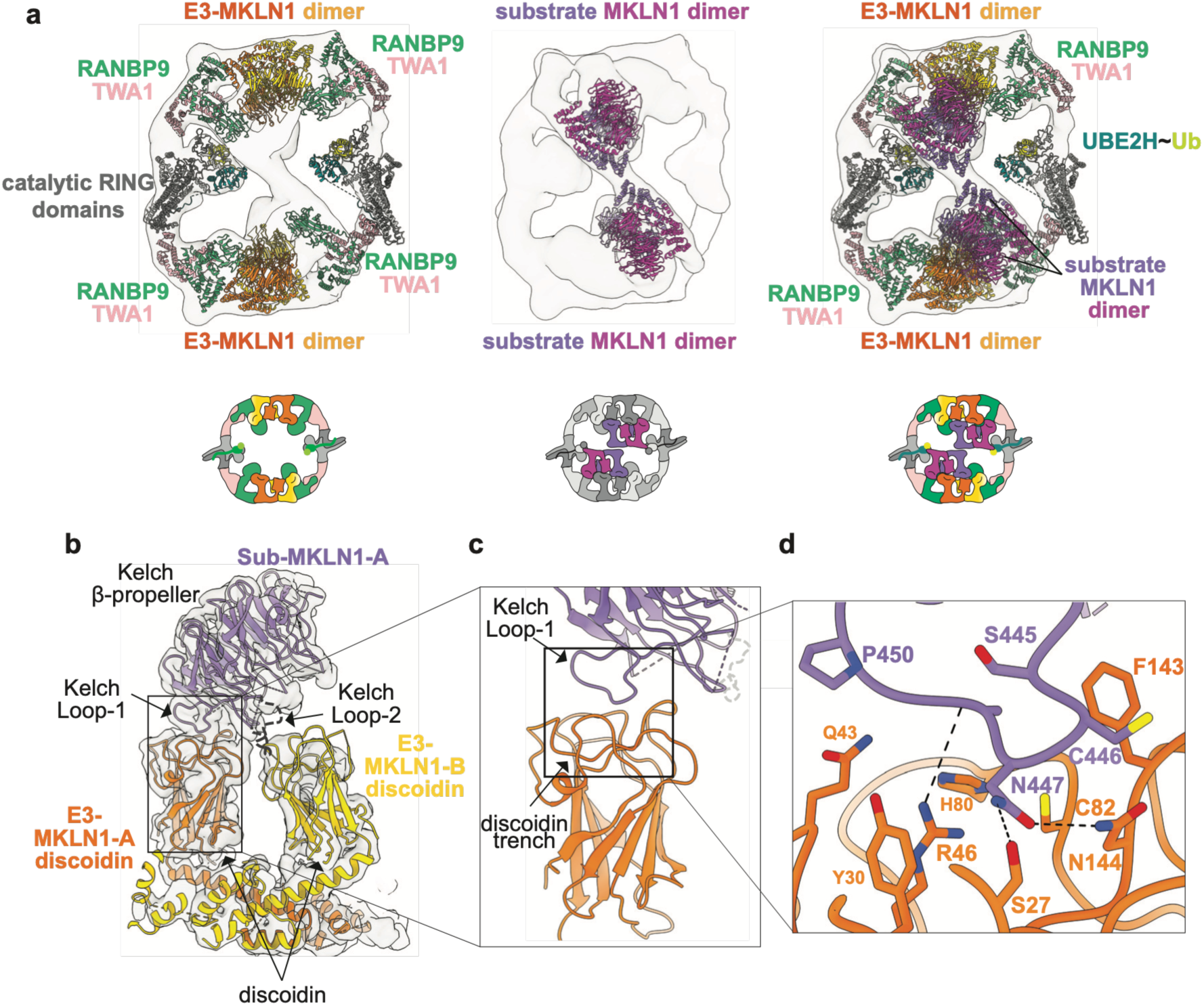
CTLH^MKLN1^-UBE2H∼ ubiquitin recruits a dimeric MKLN1 substrate through discoidin–Kelch domain contacts. **a** Cryo-EM map of CTLH^MKLN1^–UBE2H∼ ubiquitin bound to substrate MKLN1, shown with fitted structures and schematic diagrams below illustrating subunits within the assembly. Left: Cryo-EM map fitted with the scaffolding module RANBP9– TWA1 (this study), E3-MKLN1 dimer (this study), and catalytic module extracted from PDB: 8PJN. Middle: the same cryo-EM map with the MKLN1 fitted into the density corresponding to Sub-MKLN1dimer. Right: all fitted structures shown together, depicting the complete CTLH^MKLN1^-Sub-MKLN1 assembly. **b** Map of E3-MKLN1 discoidin domains engaging the Sub-MKLN1-A Kelch β-propeller resolved to 3.7 Å. Kelch Loop-1 and Loop-2 are indicated. **c** Close-up view of the boxed region in **b** showing E3-MKLN1-A discoidin trench surface and Kelch Loop-1 from Sub-MKLN1-A engaging the trench. **d** Molecular details of the E3-MKLN1-A discoidin trench– Sub-MKLN1-A Kelch Loop-1 interface, highlighting key discoidin residues contacting Sub-MKLN1-A Asn447 and surrounding loop residues, with hydrogen bonds depicted as dashed lines.

We refer to the two protomers in each MKLN1 dimer as A and B. The E3-substrate interactions were visualized by determining the structure of an E3-MKLN1-A complex with Sub-MKLN1-A from a 3.7 Å resolution focus-refined map, and then fitting domains from this structure into the E3-MKLN1-B and Sub-MKLN1-B densities in a 4.0 Å resolution focused-refined map showing the MKLN1 tetramer.

The E3-MKLN1 and substrate MKLN1 dimers are related by approximate 2-fold symmetry, such that the E3-MKLN1-A subunit is equivalent to the Sub-MKLN1-A subunit, and the E3-MKLN1-B subunit is equivalent to the Sub-MKLN1-B subunit (Supplementary Fig. 2b). Within each dimer, however, the two protomers are asymmetric. Such arrangements had been observed in prior cryo-EM maps of MKLN1 alone and in subcomplexes^15,54^, although this arrangement had not been previously observed in the context of a CTLH E3 complex. Moreover, the role of the tetrameric assembly in defining MKLN1 as a CTLH E3 substrate was not previously known.

The MKLN1 substrate is recruited to the CTLH^MKLN1^ E3 through multiple interactions. The discoidin domains from both E3 protomers simultaneously engage the Kelch propeller domain of Sub-MKLN1-A (Fig. 3b), while the Kelch domain of E3-MKLN1-A reciprocally contacts the discoidin domains of both substrate protomers (Supplementary Fig. 2c). The discoidin domain comprises an 8-stranded β-barrel with a helical turn between strands 1 and 2 (Fig. 3c).

Extended loops connecting the strands form a trench-like surface at the Kelch-facing end of the barrel. The E3-MKLN1-A discoidin domain trench embraces a five-residue loop (hereafter referred to as Loop-1) between blades 4 and 5 from the Sub-MKLN1-A Kelch domain. The Loop-1 residue Asn447 lies at the center of this interface. It embeds in the discoidin trench of E3-MKLN1-A, where it is surrounded by Tyr30, Arg46, Cys82, Asn144 and Ser27, and forms hydrogen bonds with the latter two side-chains (Fig. 3d). The remainder of the substrate loop is secured by backbone and van der Waals contacts with Gln43, His80, and Phe143 from the E3-MKLN1-A discoidin domain.

The E3-MKLN1-A discoidin domain interaction with the Sub-MKLN1-B Kelch domain is visible only at lower resolution. Nonetheless, the data show this involves a distinct discoidin domain surface contacting a “Loop-2” between the central two strands in blade 4 of the Kelch domain (Supplementary Fig. 2c).

The Kelch domains of E3-MKLN1-B and Sub-MKLN1-B are located at the distal ends of the tetrameric assembly, and are not well-resolved in our higher-resolution maps. Their positions suggest potential for interactions between E3-MKLN1-B and Sub-MKLN1-A, and between E3-MKLN1-A and Sub-MKLN1-B Kelch domains (Supplementary Fig. 2d). However, we did not obtain a map at sufficiently high resolution to allow definitive residue assignments for such interactions.

### Substrate: ZMYND19

The maps of the CTLH^MKLN1^-UBE2H∼ubiquitin complex with ZMYND19 revealed three striking features (Fig. 2a, middle). First, the only MKLN1 dimer in this map was the E3-MKLN1, i.e. within the oval CTLH E3 assembly. Second, the E3-MKLN1 dimer connects to additional density corresponding to two molecules of ZMYND19 in asymmetric arrangement. Third, different 3D classes showed varying extents of density for ZMYND19 indicating that one molecule (hereafter termed ZMYND19-A) binds stably while the second (hereafter termed ZMYND19-B) associates more dynamically (Supplementary Fig. 3a).

Maps from a MKLN1–RANBP9–TWA1 subcomplex showed detailed interactions with ZMYND19 (Fig. 4a, Supplementary Fig. 3b). For ZMYND19-A - i.e. the better-resolved molecule, one loop (hereafter 40s-loop) from the globular ZMYND domain binds the E3-MKLN1 protomer A discoidin domain trench. Another ZMYND19 loop (150s-loop) extends towards the discoidin domain trench from the E3-MKLN1-B subunit. Meanwhile, the ZMYND19 C-terminal tails from both protomers extend towards the E3-MKLN1 Kelch domains.

**Fig. 4.**
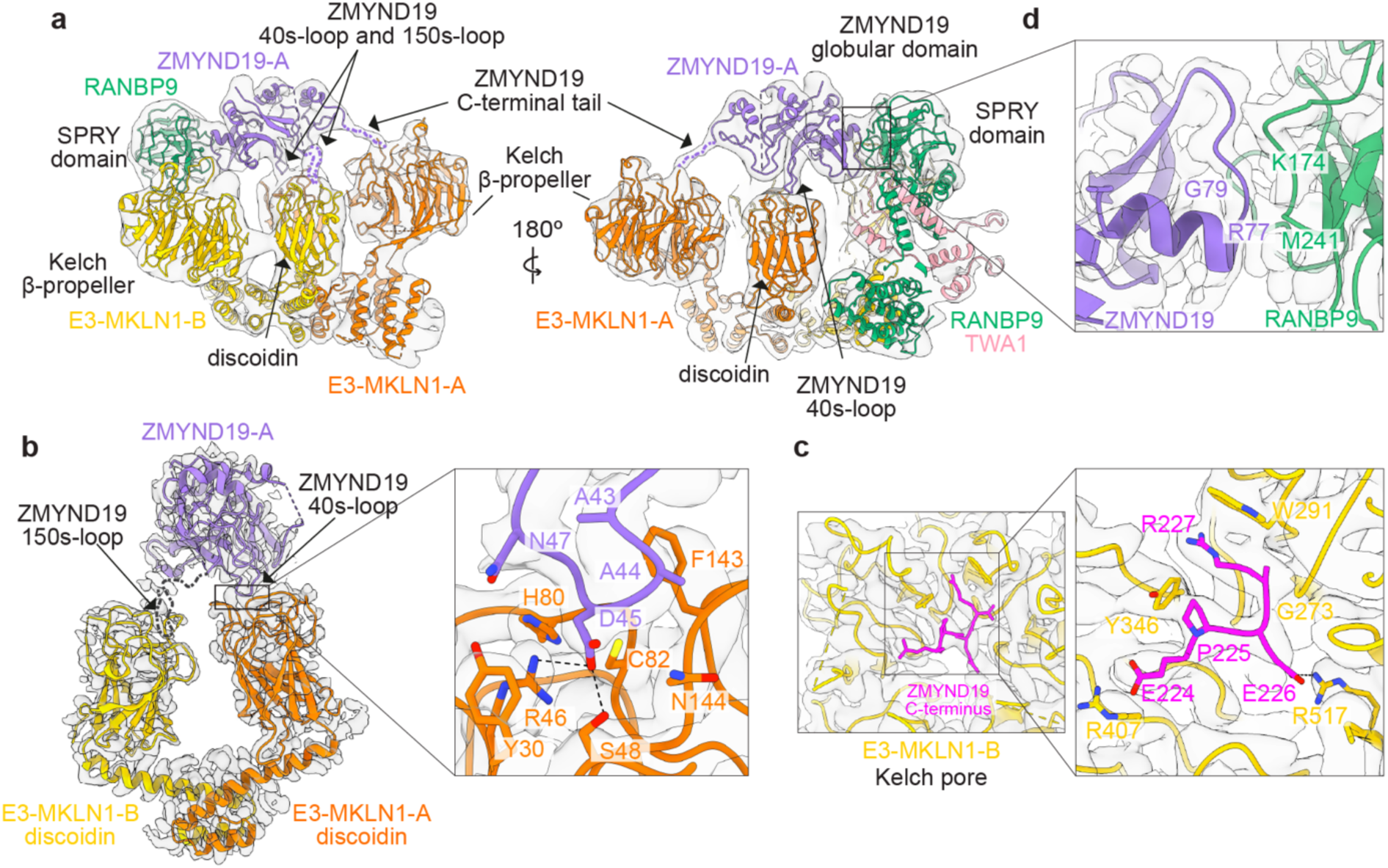
ZMYND19 is recruited to CTLH^MKLN1^ through multivalent contacts with the discoidin domain and Kelch β-propeller pore. **a** Two views of the cryo-EM map of the MKLN1–RANBP9–TWA1 subcomplex bound to ZMYND19-A, with fitted model from this study. Views illustrate ZMYND19-A loop-1 and loop-2 contacts with the E3-MKLN1-A and E3-MKLN1-B discoidin domains, the ZMYND19 C-terminal tail extending toward the E3-MKLN1-B Kelch β-propeller, and the ZMYND19-A globular domain contacts with the RANBP9 SPRY domain. Subunits and domains are indicated. **b** Focus-refined cryo-EM map resolved to 3.23 Å, with fitted model, showing ZMYND19 40s-loop engaging the E3-MKLN1-A discoidin trench. Close-up highlights molecular details of the interface, with key discoidin residues contacting ZMYND19 Asp45 and surrounding loop residues, with hydrogen bonds shown as dashed lines. **c** Focus-refined cryo-EM map resolved to 3.34 Å, showing the ZMYND19-B C-terminal residues fitted into the E3-MKLN1-B Kelch domain channel. Close-up highlights molecular details of the Arg/C-degron recognition interface, shown in sticks and hydrogen bonds depicted as dashed lines. **d** Focus-refined cryo-EM map resolved to 3.16 Å, showing backbone-mediated contacts between ZMYND19-A and the RANBP9 SPRY domain. b-d were sharpened with DeepEMhancer.

The contacts between the E3-MKLN1-A-discoidin domain and the ZMYND19 40s-loop were resolved in detail after focus-refinement (Fig 4b). Interactions center around ZMYND19 Asp45, which is surrounded by Tyr30, Arg46, Ser48, Cys82, and Asn144 from the discoidin domain trench. While these interactions resemble those of Sub-MKLN1-A Asn447, the ZMYND19 Asp45 side-chain forms electrostatic interactions with E3-MKLN1-A Arg46 and Ser48 (Fig 3d). Although limited resolution precludes unambiguous assignment of contacts with the ZMYND19 150s-loop, we note that this loop sequence (Asn153-Gly154-Asp155) is palindromic to that of the 40s-loop (Asp45-Gly46-Asn47), raising the possibility of analogous interactions in the reverse orientation.

A focus-refined map visualized the last four ZMYND19-B residues (Glu224-Pro225-Glu226-Arg227) fitting inside the E3-MKLN1-B Kelch domain central channel (Fig 4c). The terminal ZMYND19 Arg227 forms π-stacking interactions with MKLN1 Trp291. Adjacent contacts further support this Arg/C-degron. ZMYND19 Glu226 hydrogen bonds with MKLN1 Arg517. ZMYND19 Pro225 packs against the aromatic ring of MKLN1 Tyr346, and ZMYND19 Glu224 is positioned near MKLN1 Arg407.

Yet, another ZMYND domain surface of molecule A, centered around Gly78 and Gly79, approaches the SPRY domain from the RANBP9 subunit adjacent to E3-MKLN1-B (Fig 4d). These contacts are backbone-mediated, suggesting a structural rather than side-chain specific contribution to ZMYND19 recognition.

### Substrate: UNG2

Previous studies indicated a distinct substrate-binding mode for UNG2: CTLH^MKLN1^ binds FAM72A, which in turn recruits UNG2^54,61,75^. Although published cryo-EM maps for the CTLH^MKLN1^-FAM72A-UNG2 complex^54^ are consistent with this model, they do not reveal molecular details. We obtained further structural insights into this E3-substrate assembly from cryo-EM data for a complex poised for ubiquitylation, and for a subcomplex. In agreement with prior studies, the FAM72A structure consists of two domains - a globular region with a Yippee-domain fold stabilized by two zinc atoms, and a C-terminal tail. Also as described^54^, two FAM72A molecules are symmetrically positioned in the center of the oval-shaped CTLH^MKLN1^ complex. The Yippee domain of one FAM72A molecule binds the E3-MKLN1-B protomer discoidin domain, while the FAM72A C-terminus extends toward the E3-MKLN1-A Kelch domain (Fig 5a). The second FAM72A molecule makes the equivalent interactions with the opposite MKLN1 protomers.

**Fig. 5.**
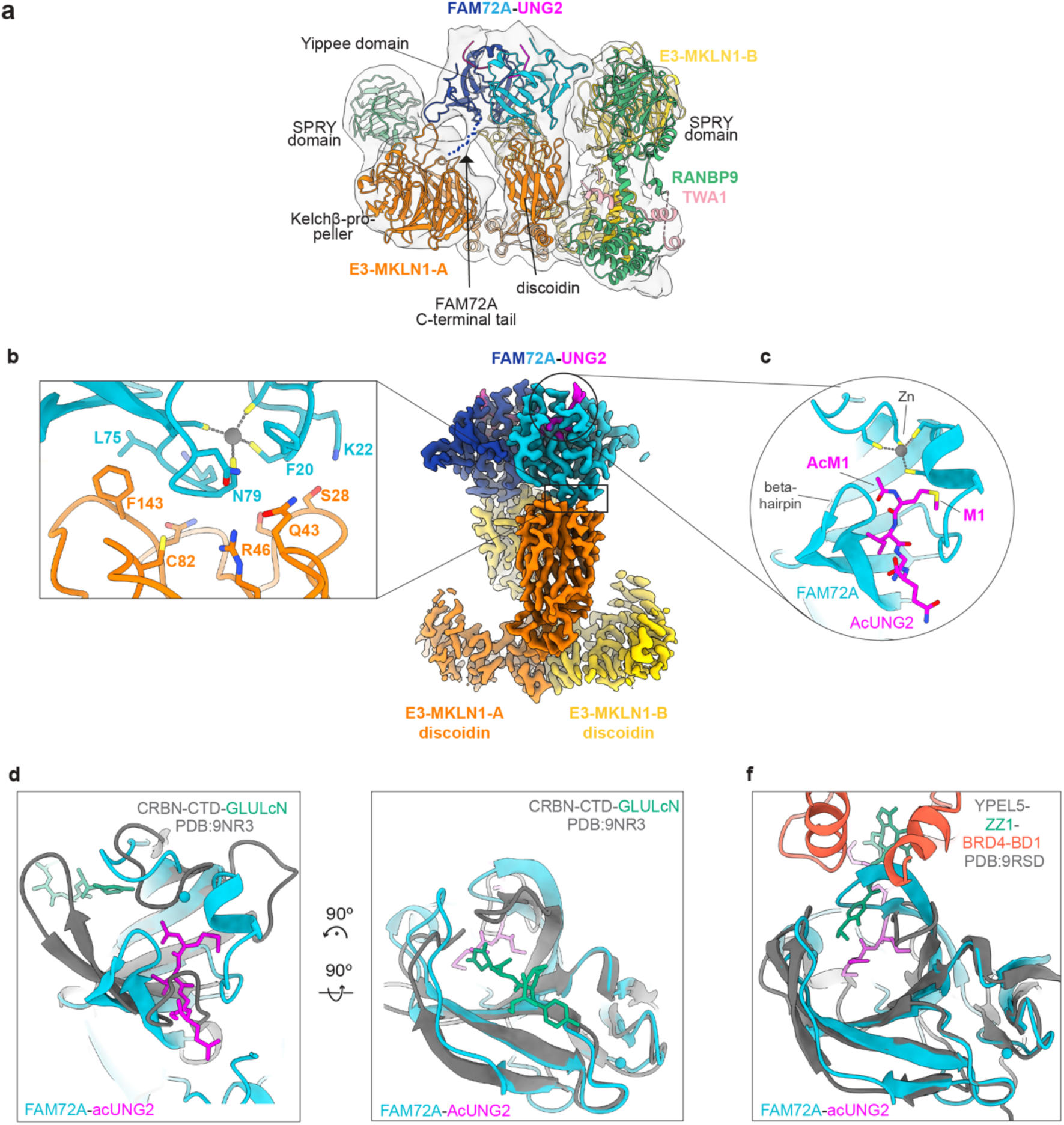
Structure of MKLN1-FAM72A–UNG2 reveals FAM72A Yippee domain engagement of the discoidin trench and FAM72A-mediated recognition of the acetylated UNG2 N-terminus. **a** Cryo-EM map of the MKLN1-RANBP9-TWA1 subcomplex bound to FAM72A-UNG2, fit with structures from this study. The composite model shows each FAM72A engaging the MKLN1 discoidin domain, and the FAM72A C-terminal tail extending toward the opposite MKLN1 Kelch β-propeller. **b** Left: close-up of the FAM72A zinc-coordinated Yippee fold–MKLN1 discoidin trench interface, with key interacting residues labelled. Middle: 3.0 Å resolution DeepEmhancer-sharpened cryo-EM map of the MKLN1 discoidin-FAM72A–UNG2. **c** Close-up of the UNG2 acetylated N-terminus (AcM1, magenta) inserted into the Y-junction tunnel inside the FAM72A Yippee domain. **d** Structural comparison of FAM72A–AcUNG2 (cyan) with C-terminal cyclic imide-bound CRBN_CTD_ (PDB 9NR3), illustrating both complexes recognizing protein termini at the center of the groove despite opposite substrate orientations. **e** Structural comparison of FAM72A–AcUNG2 with YPEL5-ZZ1-BRD4_BD1_ (PDB 9RSD), showing similar substrate positioning relative to the Yippee domain groove. The FAM72A-specific zinc-binding element and extended β-hairpin enclosing the UNG2 acetylated N-terminus are indicated.

Our 3.0 Å resolution map uniquely resolved the detailed interactions between the MKLN1 discoidin domain and FAM72A Yippee domain, and the basis for UNG2 recruitment to FAM72A (Fig 5b). A convex zinc-coordinated portion of the FAM72A Yippee fold complements the concave surface of the MKLN1 discoidin trench. A FAM72A loop - adjacent to the zinc-binding site - embeds in the trench.

The structure also shows UNG2’s acetylated N-terminal region inserted into tunnels arranged like a Y-junction inside FAM72A (Fig. 5c). UNG2 residues 2-5 insert through a long tunnel. The intersecting tunnels are punctuated by the second zinc-binding site, and accommodate the acetylated N-terminal Met1. The acetyl and Met1 side-chain are splayed apart, and directed towards opposite sides of the Y-junction. Thus, our cryo-EM data now provide a structural understanding for many previous biochemical findings indicating that the N-terminal region of UNG2 is crucial for binding FAM72A^54,76,77^.

FAM72A’s substrate-binding domain shares a common fold with CRBN and YPEL5. Comparing their modes of substrate recruitment shows their distinguishing features. Superimposing FAM72A and CRBN shows that they engage targets in inverse directions, at opposite ends of their substrate-binding grooves^78^. Yet, they both recognize protein termini - an acetylated N-terminus or C-terminal cyclic imide - in the middle of the groove (Fig. 5d). FAM72A and YPEL5 (together with a degrader molecule) recruit their targets to the same side of the groove, but YPEL5 lacks the FAM72A-specific zinc-binding element that binds UNG2’s acetylated Met1 (Fig. 5e)^48,51^.FAM72A is uniquely closed around its substrate: an extended FAM72A beta-hairpin wraps around UNG2 to approach the second zinc-binding region in FAM72A.

## Effects of mutations in MKLN1 surfaces

### MKLN1 Kelch domain

Altogether, the cryo-EM structures showed two distinct regions of the MKLN1 Kelch domain involved in substrate or adaptor recruitment. First, the MKLN1 substrate is recruited via Kelch domain loops. A Loop-1 N447R substitution designed to disrupt interaction in trans with another MKLN1’s discoidin domain converted isolated MKLN1 from a tetramer to a dimer (Supplementary Fig. 4a). This mutation also eliminated MKLN1 ubiquitylation in vitro, and stabilized MKLN1 levels in cells (Fig. 6a-b). Similar effects were observed in the in vitro assays upon wholesale replacement of the Loop-1 sequence (Supplementary Fig. 4a, b). Strikingly, the N447R mutation enhanced ubiquitylation of ZMYND19 and UNG2 (Fig. 6a). This effect agrees with our structural data showing that Sub-MKLN1 binding is mutually exclusive with other substrates, and that neither ZMYND19 nor FAM72A engages the Kelch Loop-1 surface. The data indicate that MKLN1 self-recruitment limits CTLH^MKLN1^ E3 access to other substrates competing for the discoidin domain trench.

**Fig. 6.**
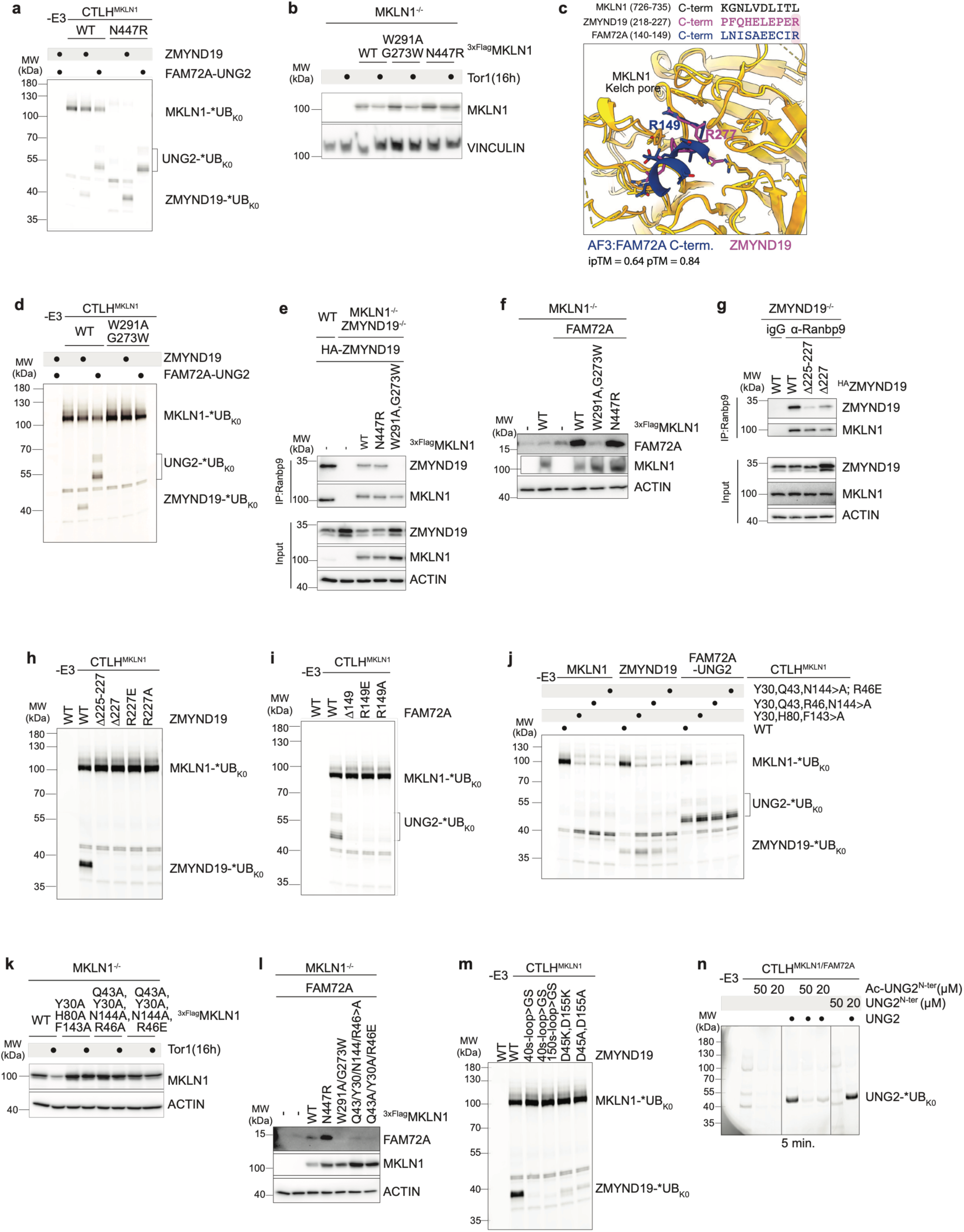
Structure-guided mutational analysis of CTLH^MKLN1^ substrate recognition. Unless otherwise noted, in vitro ubiquitylation assays were performed for 15 min using N-terminally fluorescently labeled K0-ubiquitin (*UB_K0_) with UBE2H as the E2. All cellular experiments were performed in HEK293 cells. Assays are representative of n = 2 technically independent experiments. **a** In vitro ubiquitylation assay comparing CTLH^MKLN1^ WT and the N447R mutant toward MKLN1, ZMYND19, and FAM72A-UNG2. **b** Immunoblot analysis of MKLN1 levels in MKLN1-deficient cells transfected with WT, W291A/G273W, or N447R MKLN1 following mock or torin1 treatment (16h). Vinculin serves as loading control. **c** Top: sequence alignment of MKLN1, ZMYND19, and FAM72A C-termini. Bottom: AlphaFold3 model (ipTM = 0.64, pTM = 0.84) of FAM72A C-terminus docked in the MKLN1 Kelch central channel superimposed on the cryo-EM structure of ZMYND19 C-terminal residues (this study), with terminal Arg residues R149 (FAM72A) and R277 (ZMYND19) indicated. **d** In vitro ubiquitylation assay examining the effect of the W291A/G273W Kelch central channel mutant on CTLH^MKLN1^ activity toward MKLN1, ZMYND19 and FAM72A-UNG2. **e** Co-immunoprecipitation via anti-RANBP9 antibody in MKLN1 and ZMYND19-deficient HEK293 cells co-transfected with HA-ZMYND19 and 3×Flag-MKLN1 WT, N447R, or W291A/G273W mutants, assessing the effect of Kelch channel mutations on ZMYND19 association with the CTLH complex. Immunoprecipitates and inputs are shown. Actin serves as loading control. **f** Immunoblot of FAM72A and MKLN1 levels in MKLN1 deficient cells co-transfected with untagged FAM72A and 3×Flag-MKLN1 WT, W291A/G273W, or N447R. Actin serves as loading control. **g** As in **e**, but in ZMYND19 deficient HEK293 cells transfected with HA-ZMYND19 WT, Δ225-227, or ΔR227, assessing ZMYND19 C-terminal mutant association with the CTLH complex and rabbit IgG as a negative control. **h-i** In vitro ubiquitylation assays examining the effect of C-terminal mutations in ZMYND19 (**h**) or FAM72A (**i**) on ubiquitylation by CTLH^MKLN1^. **j** In vitro ubiquitylation assays comparing CTLH^MKLN1^ WT and discoidin trench mutants toward ZMYND19, FAM72A-UNG2, and MKLN1. **k** Immunoblot of MKLN1 levels in MKLN1 deficient cells transfected with 3×Flag-MKLN1 WT or discoidin trench, following mock or torin1 treatment (16h). Actin serves as loading control. **l** Immunoblot of FAM72A and MKLN1 levels in MKLN1 deficient cells, co-transfected with untagged FAM72A and 3×Flag-MKLN1 WT or discoidin trench mutants. Actin serves as loading control. **m** In vitro ubiquitylation assays examining the effect of ZMYND19 trench-inserting loop substitutions (40s-loop>GS; 40s-loop and 150s-loop>GS) and central loop residue substitutions (D45K/D155K; D45A/D155A) on ZMYND19 ubiquitylation by CTLH^MKLN1^. **n** In vitro ubiquitylation assays testing competition of acetylated or non-acetylated UNG2 N-terminal peptides (corresponding to the first 25 amino acids of UNG2) with full-length UNG2 with for CTLH^MKLN1^-FAM72A E3 (complex co-expressed as a single construct; see Materials and Methods). These reactions were performed for 5 min.

**Table 1.**
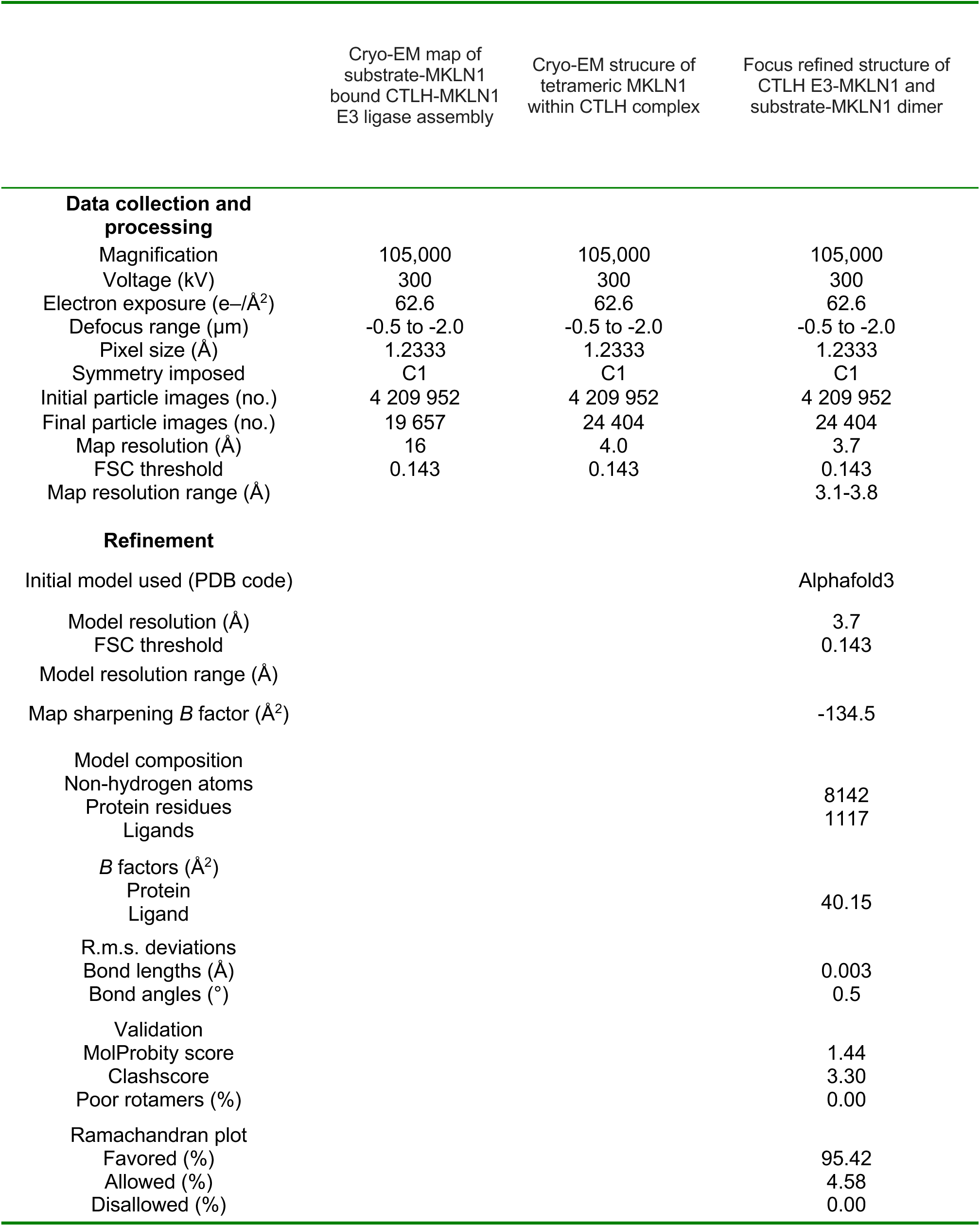
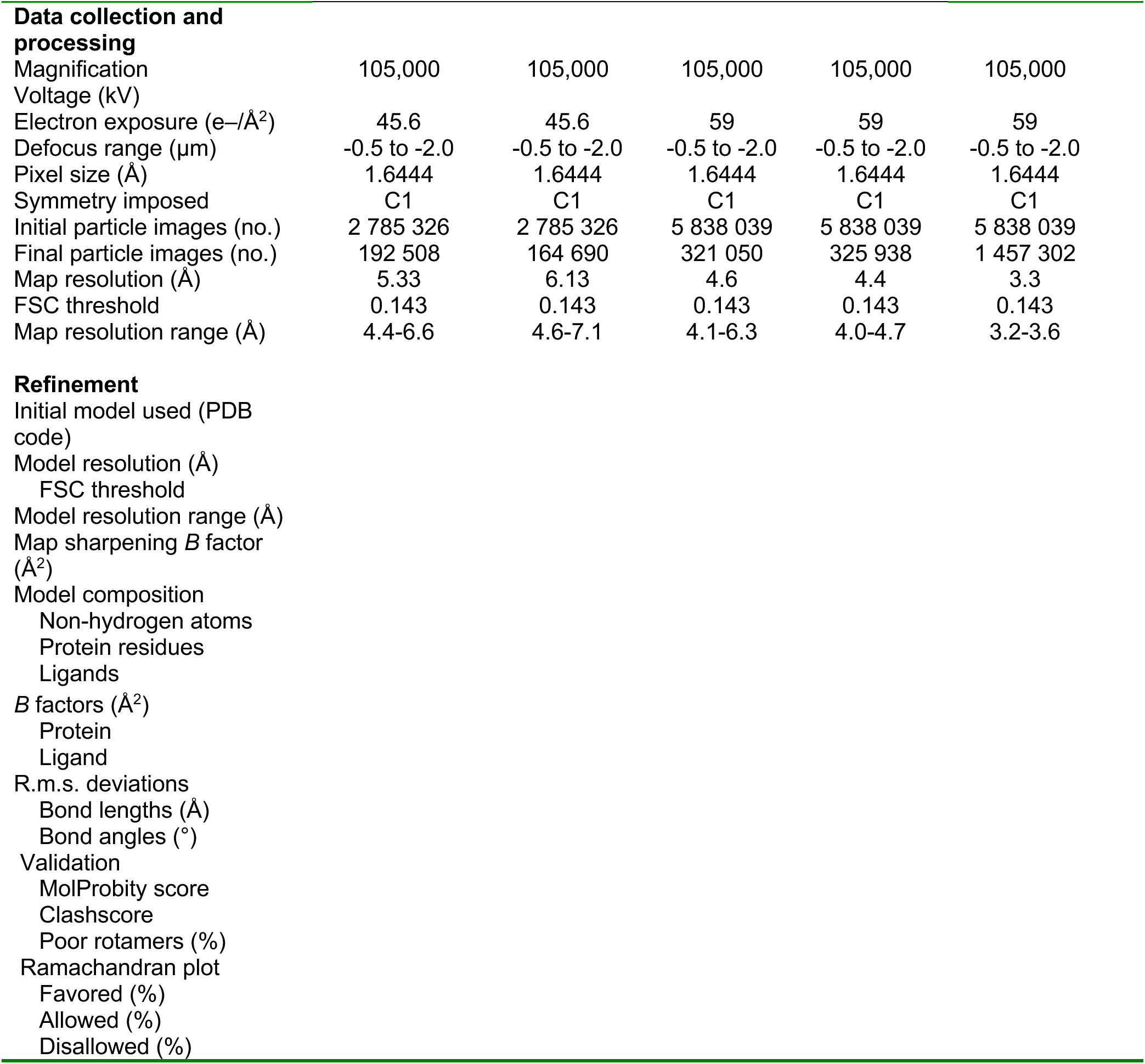
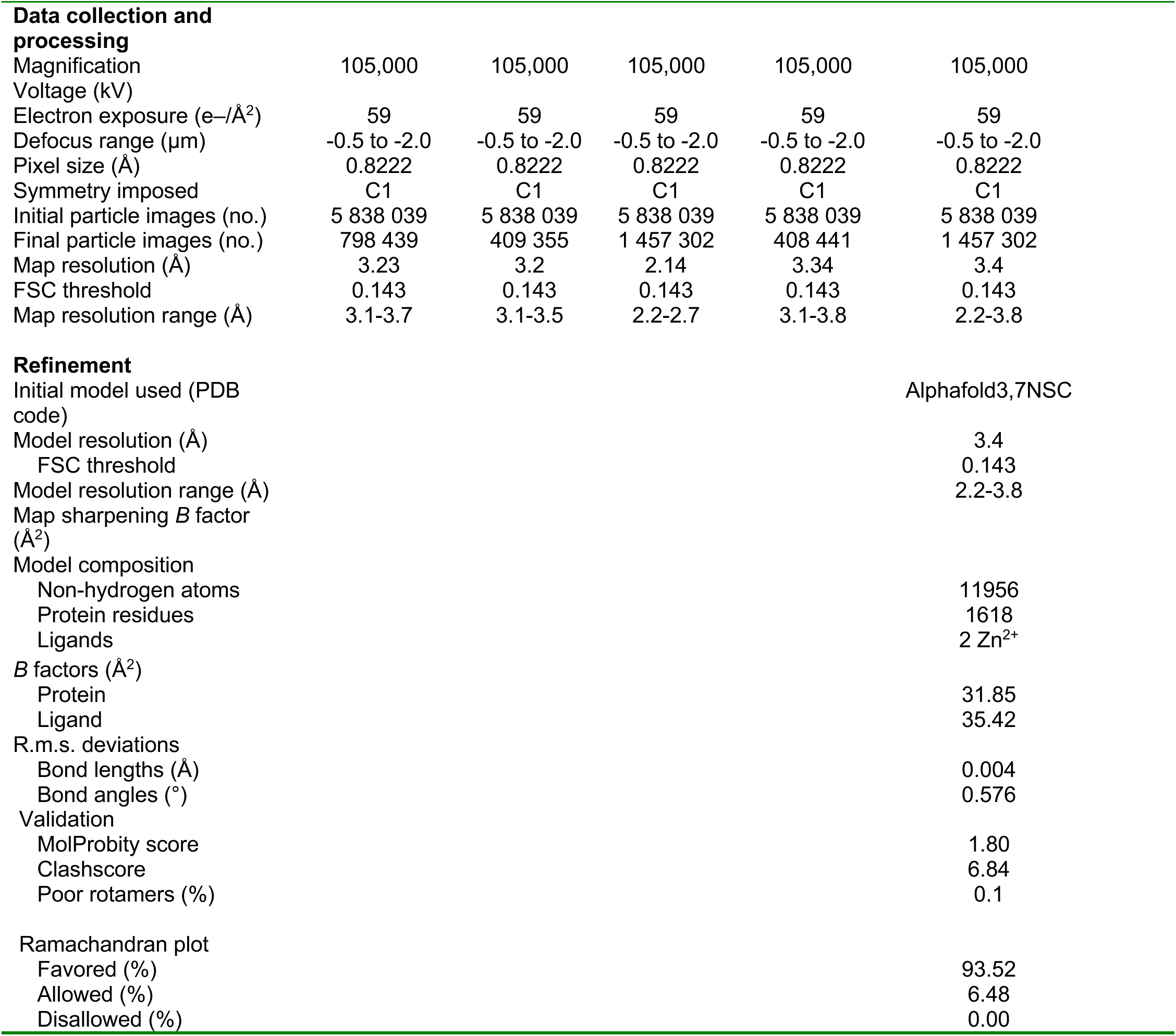
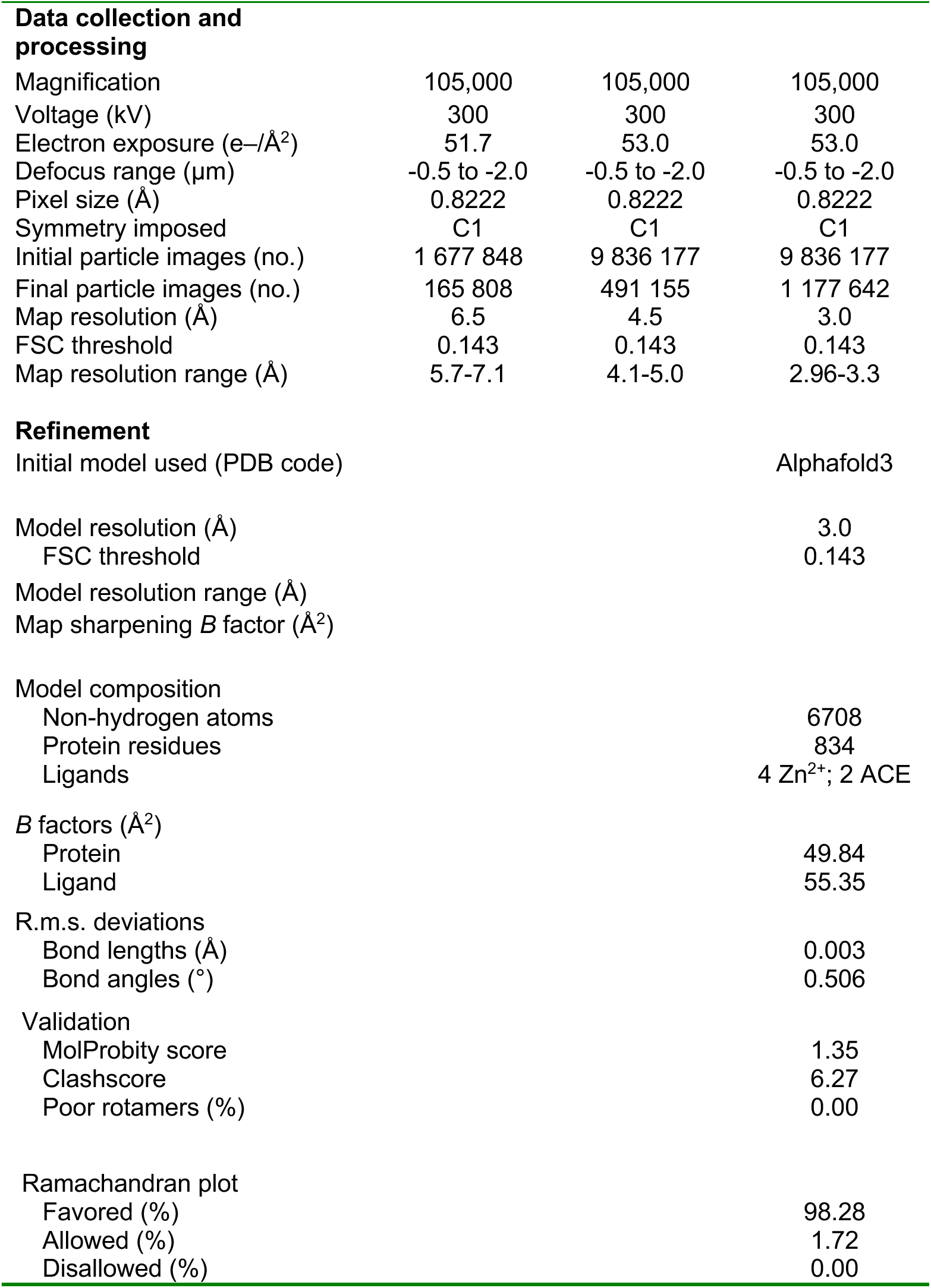
Data collection, refinement and validation statistics.

The second role for the E3-MKLN1 Kelch domain is its central channel binding a C-terminal Arg, as observed at high-resolution in our structure with ZMYND19. Also, lower resolution cryo-EM data showed the C-terminal tail of FAM72A poised to make a similar interaction (Fig. 4c, 5a). Sequence comparison confirms that ZMYND19 and FAM72A both terminate with an Arg, whereas MKLN1 does not. AlphaFold3 predicts that the FAM72A C-terminus binds the MKLN1 Kelch domain channel much like the cryo-EM structure with ZMYND19 (Fig. 6c)^79^. Accordingly, a MKLN1 mutant with residue substitutions in the Kelch central channel (W291A and G273W) was impaired for in vitro ubiquitylation of ZMYND19 and the FAM72A-dependent substrate UNG2 (Fig. 6d). Upon co-expression in HEK293 cells, the association of ZMYND19 with the CTLH complex was disrupted for the Kelch channel mutant (W291A and G273W mutant) (Fig 6e). Meanwhile, the levels of exogenously expressed FAM72A substantially increased upon co-expression with MKLN1, but not with the W291A and G273W mutant (Fig. 6f). The latter data suggest that MKLN1 may protect FAM72A from degradation, potentially through promoting its folding, much like levels of the structurally homologous YPEL5 depend on co-expression of its Gid7-like partner WDR26^48^. Notably, mutation of the Kelch channel did not affect MKLN1 ubiquitylation or degradation (Fig. 6b, d), consistent with our structural data showing Sub-MKLN1 does not engage through this region.

Several experiments considering MKLN1 Kelch channel engagement from the perspective of its binding partners showed a key role for their C-terminal Arg. In cellular assays, C-terminal mutants in ZMYND19 were impaired for association with the CTLH complex (Fig. 6g). Moreover, ZMYND19 ubiquitylation in vitro was essentially abolished for C-terminal deletion mutants and the C-terminal R227E mutant. Shortening the C-terminal side-chain with an R227A substitution partially reduced ubiquitylation with purified proteins (Fig. 6h). Likewise, deleting the FAM72A C-terminal Arg or mutating it to Asp or Ala also impaired UNG2 ubiquitylation (Fig. 6i). Thus, the structural and biochemical data nominate MKLN1 as an Arg/C-degron-binding E3. This concept was also proposed by work posted on bioRxiv while our manuscript was in preparation^67,68^.

### MKLN1 Discoidin domain

The structural data show that a plethora of side-chains from the MKLN1 discoidin domain trench differentially contact various MKLN1 binding partners. Ala replacements for MKLN1 discoidin domain residues Tyr30, His80, and Phe143 - which contact the opposing MKLN1 Kelch domain Loop-1 (Fig. 3d) - reduced MKLN1 ubiquitylation in vitro (Fig. 6j). Importantly, the effects of this discoidin domain mutant paralleled those of the Kelch Loop-1 mutant (see Fig. 6a, Supplementary Fig. 4b), with increased alternative autoubiquitylation of other CTLH E3 subunits, and slightly increased ubiquitylation of ZMYND19 and UNG2 (Fig. 6j). Meanwhile, eliminating a different constellation of side-chains that also include a key ZMYND19-binding residue (R46E) reduced ubiquitylation of ZMYND19 along with MKLN1. In cells, these mutants block the MKLN1 degradation pathway, and lose ability to support FAM72A levels (Fig. 6k, 6l).

In vitro ubiquitylation of ZMYND19 was also impaired by mutation of the ZMYND domain loops that bind the MKLN1 discoidin domains (40s-loop alone, or together with mutation of the 150s-loop, Fig. 6m). Similar results were also observed for mutation of the central Asp residues from the two loops (Asp45 from the 40s-Loop and Asp155 from the 150s-Loop). However, discoidin domain mutants at the interface with ZMYND19, and these ZMYND19 loop mutants retained interaction when exogenously expressed in cells (Supplementary Fig. 4c and d). Thus, under the conditions of our cellular assays, the ZMYND19 interactions with the MKLN1 Kelch domain, or other regions of the CTLH E3, may be sufficient to mediate interaction.

### Role of UNG2 N-terminal acetylation in binding FAM72A

The CTLH^MKLN1^-FAM72A recognition of UNG2’s acetylated N-terminus was striking considering that the first-characterized substrates of GID/CTLH E3s are recognized by N-degrons binding to the GID4 subunit^5,39^. To determine the importance of the acetyl group for FAM72A binding, we tested whether peptides corresponding to the UNG2 N-terminal region could compete with full-length substrate^80^. Addition of the N-terminally acetylated peptide reduced UNG2 ubiquitylation compared to the non-acetylated counterpart (Fig. 6n).

## Discussion

In determining cryo-EM structures representing three distinct CTLH^MKLN1^ ubiquitylation complexes, our data unveil how the combination of multiple domains from an E3 ligase receptor subunit allow a tremendous versatility of routes for substrate targeting. Despite unique features of three different substrates, they all converge on two common themes: multivalent engagement of the E3 ligase - either by the substrate itself or a substrate-binding adaptor, and placement near the ubiquitylation active sites within a giant oval E3 ligase assembly (Fig. 2).

Each E3-MKLN1 dimer displays four interaction domains that altogether can be engaged in various ways (Fig. 2c). The Kelch domain Loop-1 cooperates with a particular portion of the discoidin domain trench surface to recruit the Sub-MKLN1 dimer in a 2-fold symmetric arrangement (Fig. 3). Multiple discoidin and the Kelch domain surfaces can be asymmetrically engaged, as shown by their recruiting a pair of ZMYND19 molecules (Fig. 4). Meanwhile, the discoidin trench and Kelch propeller surfaces can also mediate symmetric interactions, as during engagement of a pair of FAM72A-UNG2 complexes (Fig. 5). And the UNG2 substrate is secured through its acetylated N-terminus engaging the Yippee-domain groove of FAM72A (Fig. 5c). The finding that natural substrates exploit multiple contact points distributed across MKLN1 and its binding partners has functional implications: E3 engagement at multiple sites can increase binding specificity, reinforce individually weak contacts, or constrain domain positions that would otherwise be flexibly tethered.

The structures nominate several potential MKLN1-dependent degron motifs. Internal loops from both Sub-MKLN1 and ZMYND19 display a central Asp or Asn, respectively, embedding in the MKLN1 discoidin domain trench. The second, less-resolved ZMYND19 loop that contacts the other E3 MKLN1 protomer’s discoidin also contains an Asp and an Asn residue that are candidates for making similar interactions, with potential for the loop arranged in the opposite orientation. Despite these similarities, the breadth of distinct substrate contacts with the MKLN1 discoidin domain allows design of E3 mutations selectively impairing association with subsets of substrates (Fig. 6j). It seems likely that many CTLH substrates will be discovered that exploit the versatility of side chains exposed in the MKLN1 discoidin domain. However, the precise sequence rules governing which loop-containing proteins are recognized by the discoidin trench remain to be defined.

In addition to their loops binding the discoidin domain, both ZMYND19 and FAM72A also project C-terminal sequences that end with an Arg that captures the MKLN1 Kelch domain (Fig. 6c, h, i). This C-terminal Arg-mediated interaction appears to drive affinity for FAM72A, because in vitro ubiquitylation of UNG2 is drastically reduced upon elimination of the FAM72A C-terminal Arg or its docking site in the MKLN1 Kelch domain, whereas there is little such effect for mutations in the MKLN1 discoidin domain. The C-terminal Arg motif was also implicated as a MKLN1-binding degron in parallel studies posted on bioRxiv during preparation of our manuscript^67,68^, and notably is also in the previously reported MKLN1 substrate AAMP^81^.

Yet another mode of substrate recruitment was resolved at high-resolution in our cryo-EM structure of the MKLN1-FAM72A-UNG2 subcomplex. UNG2’s acetylated N-terminus is gripped within a Y-shaped tunnel buried inside the FAM72A Yippee domain. This architecture explains prior findings that the UNG2 N-terminal region is necessary for degradation and sufficient for FAM72A binding, and is consistent with our peptide competition assay demonstrating a critical role for N-terminal acetylation^54,76^. Interestingly, the deep tunnel in FAM72A is established by loops wrapping around UNG2’s N-terminus. We speculate that these loops could adopt alternative, more open arrangements to engage other sequences, much like the Yippee domains of YPEL5 or CRBN^48,51,82,83^. It is possible that alternative FAM72A conformations could enable recruitment of other substrates to CTLH^MKLN1^. Irrespective of whether there are additional FAM72A-recruited substrates and how they are recognized, our structural data defines how an E3 ligase can target substrates through an acetylated N-terminus. Our results also extend the CTLH E3’s capacity for recognizing substrate N-degrons to a binding mode that is mechanistically distinct from the other CTLH substrate receptor, GID4, which directly contacts the charged or polar terminus of peptides or peptidic small molecules initiating with Pro or hydrophobic residues^41,44–46^.

Beyond showing modes of CTLH^MKLN1^ substrate recognition, our data reveal how MKLN1 self-assembly mediates its ubiquitylation in a manner that competes with substrate modification. Indeed, in our in vitro ubiquitylation assays, addition of substrates ZMYND19, or FAM72A-recruited UNG2, slightly reduced ubiquitylation of WT MKLN1. Meanwhile, mutations designed to selectively disrupt the E3-MKLN1–Sub-MKLN1 interface both reduced its ubiquitylation and increased the modification of substrates. (Fig. 6a, j). Thus, MKLN1 joins the growing list of multiprotein E3 ligases that form higher-order assemblies that compete with substrates^9,84–87^. For such E3s in the cullin-RING ligase (CRL) family, substrate binding leads to activation of the E3 ligase, not only by overcoming autoinhibition, but also by directing retention of the NEDD8 modification that both protects the E3 ligase complex from disassembly and stimulates ubiquitylation activity^88,89^. Based on this precedent, we speculate that future studies could reveal additional layers of CTLH E3 regulation through formation of the MKLN1 substrate complex. For example, it is possible that the MKLN1 substrate complex could serve as a chaperone function that assists or stabilizes formation of the giant oval, or that this protects MKLN1 complexes from shuffling of CTLH subunits into complexes with WDR26 instead. The MKLN1 substrate complex may serve to protect the CTLH assembly from self-modifications that could alter activity. The precise cellular signals that tip the balance between MKLN1 self-recruitment and engagement of other substrates, whether through post-translational modifications of Kelch Loop-1, changes in MKLN1 stoichiometry, or competition from binding partners, remain to be determined.

The variety of substrate recognition modes deployed by CTLH^MKLN1^ adds to an increasingly complex picture of degron recognition across the GID/CTLH family. The originally characterized Pro/N-degrons bind GID4^41^, while basic-aromatic internal degrons bind the WD40 domain of WDR26^48^. Heterobifunctional and molecular glue degraders can recruit substrates to GID4 and to YPEL5 bound to WDR26^51,90,91^. Ours and others’ studies^67,68^ of CTLH^MKLN1^ now extend this repertoire to C-terminal Arg degrons, while our studies also reveal Asp/Asn-loop internal degrons, and adaptor-mediated Ac/N-degron recognition. Perhaps the most striking feature of the CTLH E3 is that the diverse modes of substrate recruitment all allow placement of substrates within the hollow oval E3, proximal to the active sites of the CTLH-specific E2 UBE2H. This unique E3 geometry - which allows substrate receptors to be localized at various positions along the oval and yet place substrates in the center - contrasts with the more constrained placement of substrate-binding modules in the Cullin-RING ligase family^88^, yet may serve a similar function: ensuring that diverse degron-bearing substrates are reliably positioned for ubiquitylation. Such plug-and-play E3 ligase mechanisms may be particularly suitable for targeted protein degradation.

## Acknowledgements

We thank J. F. for general scientific advice and discussion, and cryo-EM processing assistance, L. S. for protein purification assistance, general discussion and support, J.R. P. and E. S. for model building help, F. C. for writing assistance, and other members of the Schulman lab, especially J. S., G. A., M. K, A. S.T., C. B., and J. K.

We also thank Daniel Bollschweiler and Tillman Schäfer at the MPIB Cryo-EM Facility (RRID:SCR_025744), Stephan Uebel and Mohammadreza Taheri at the MPIB Biochemistry Core Facility (RRID:SCR_025743), and Barbara Steigenberger and Victoria Sánchez at the MPIB Mass Spectrometry Facility (RRID:SCR_025745).

This study was co-funded by the Max Planck Society and the European Union (ERC, UPSmeetMet, 101098161, BAS). S.A.M and E.C.P. were supported by PhD fellowships from the Boehringer Ingelheim Fonds. Views and opinions expressed are however those of the authors only and do not necessarily reflect those of the European Union or the European Research Council. Neither the European Union nor the granting authority can be held responsible for them.

## Author contributions

Conception: B.A.S., S.S.; Biochemistry: S.S.; Cryo-EM, structure building and refinement: S.S., J.D., S.A.M., J.C.; Cell biology: K.V.G.; Reagent production: S.S., S.vG.; Data analysis: S.S., K.V.G., J.D., S.A.M, J.C., E.C.P., B.A.S.; Supervision: B.A.S.; Paper preparation: S.S., K.V.G., B.A.S., with input from all authors

## Competing interests

B.A.S is a member of the scientific advisory boards of Proxygen and Lyterian, and has financial interest in Nextech Invest. The other authors declare no competing interests.

---

**Supplementary Fig. 1.**
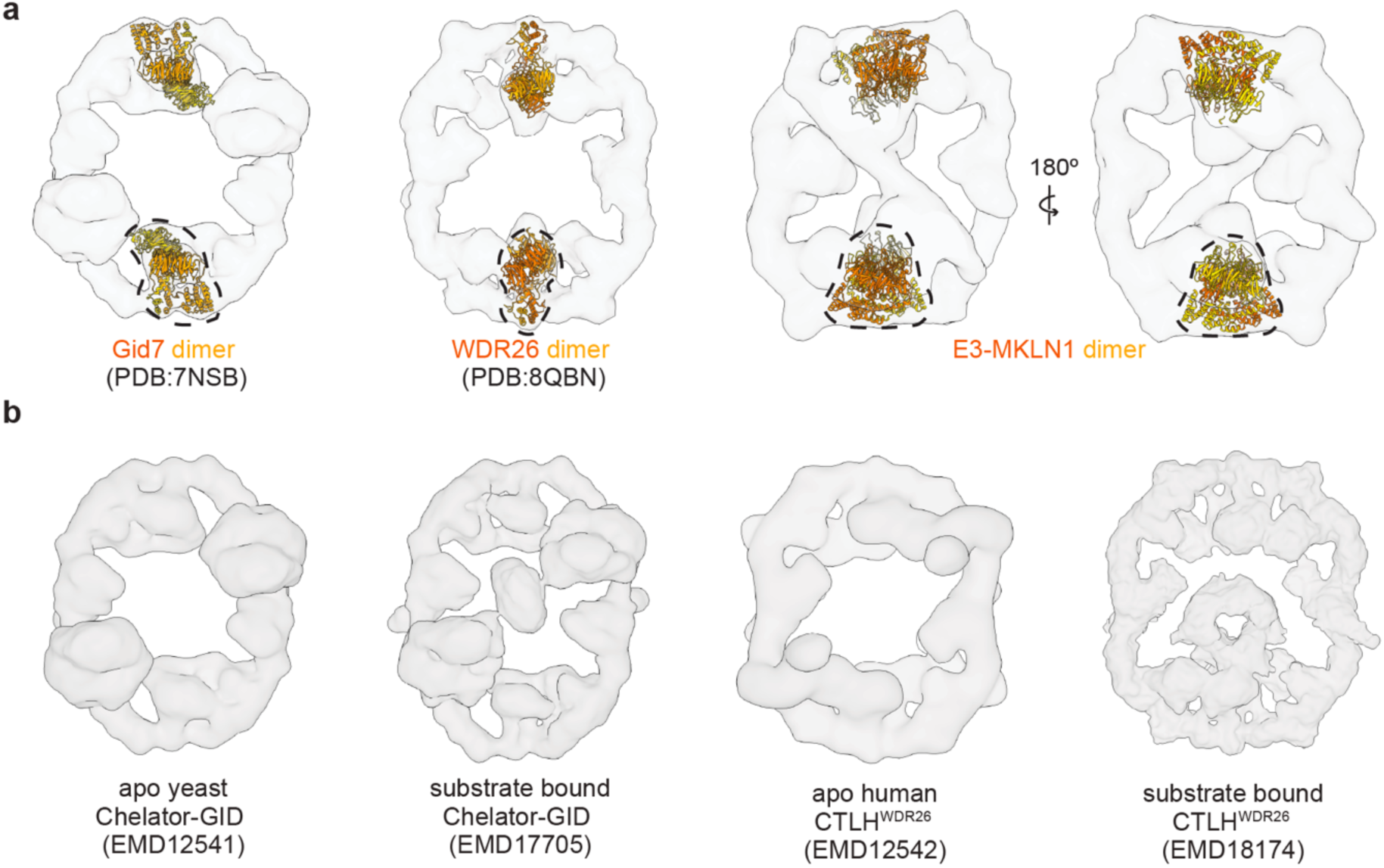
Survey of giant oval supramolecular GID/CTLH E3 assemblies. **a** Cryo-EM maps, from left-to-right, shown as transparent surface for substrate-free (apo) yeast Chelator-GID (EMD-12541) and human CTLH^WDR^^26^-YPEL5 (EMD-12542), or CTLH^MKLN1^ (this study). Dashed circles indicate the position of the Gid7-like dimer within each assembly. The indicated dimer structures of yeast Gid7 WDR26, and model for E3-MKLN1 (this study) are shown in ribbon diagram fit inside the maps (PDB: 7NSB, 8QBN, and XXXX, respectively). **b** Cryo-EM maps of substrate-free and substrate (Fbp1)-bound yeast Chelator-GID (EMD-12541 and EMD-17705), and substrate-free and substrate (NMNAT1)-bound human CTLH^WDR^^26^ (EMD-12542 and EMD-18174), illustrating how substrates are engaged inside the hollow central cavity of oval GID/CTLH E3 assemblies, much like Sub-MKLN1 fills the center of CTLH^MKLN1^ (Fig. 3a).

**Supplementary Fig. 2.**
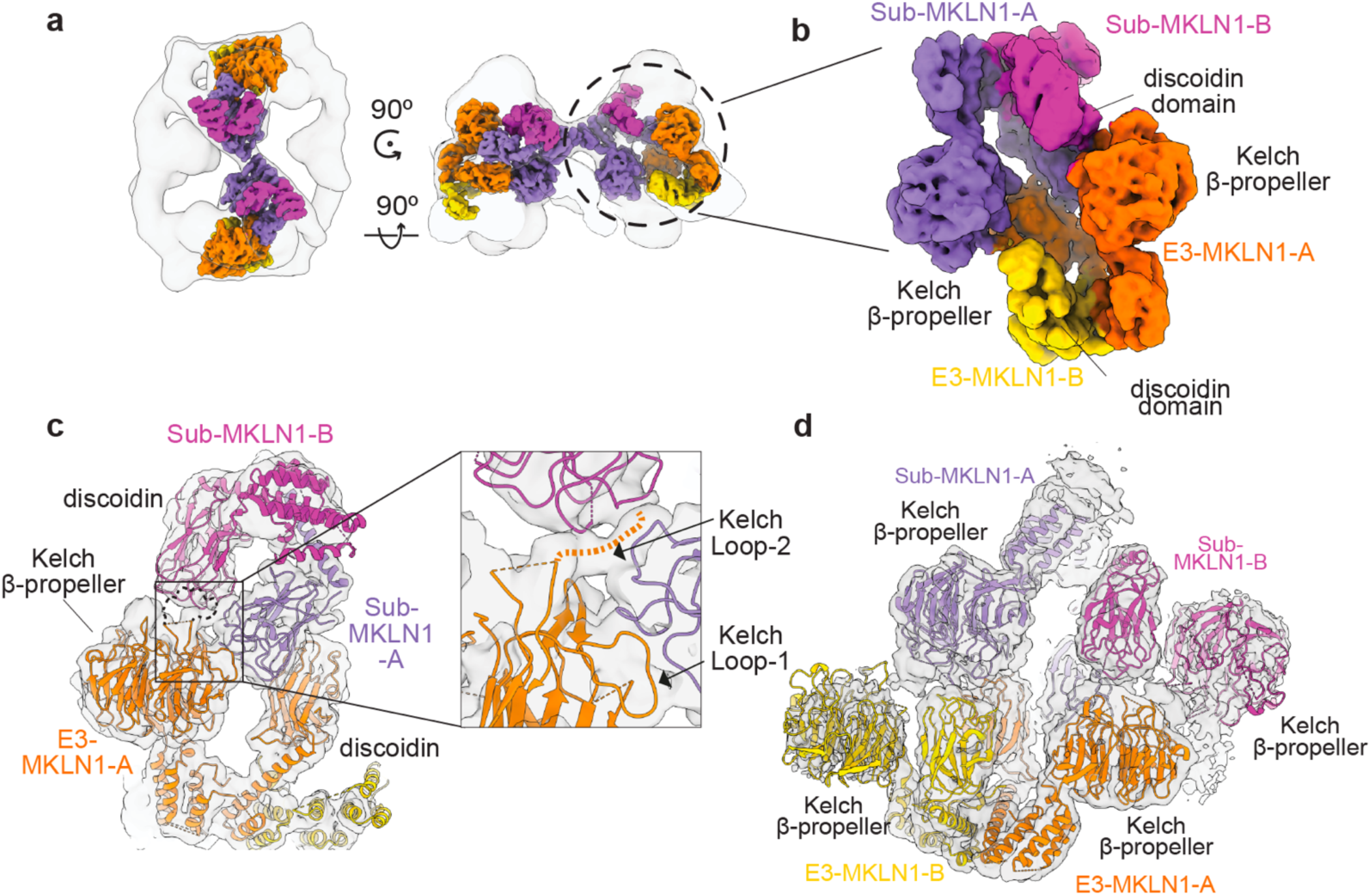
Cryo-EM analysis of the E3-MKLN1–Sub-MKLN1 tetrameric assembly. **a** Cryo-EM map of CTLH^MKLN1^-UBE2H∼ubiquitin (transparent) fit with focus-refined map of E3-MKLN1–Sub-MKLN1 tetramer assembly (solid surface colored by MKLN1 protomer). **b** Close-up view of cryo-EM density of tetrameric MKLN1 arrangement illustrating the approximate 2-fold symmetry relating the E3-MKLN1 and Sub-MKLN1 dimers. Within each dimer, the MKLN1 protomers are asymmetrically arranged. **c** Portion of focus-refined map of E3-MKLN1–Sub-MKLN1 tetramer assembly shown as a transparent surface, fitted with model derived from E3-MKLN1–Sub-MKLN1 dimer structure. One protomer E3-MKLN1-A (dark orange) was fit from high-resolution structure of its complex with Sub-MKLN1 (PDB: XXXX). The other protomer (E3-MKLN1-B, yellow) was modeled by fitting the domains (discoidin, LisH-CTLH-CRA, Kelch) from the E3-MKLN1-A structure individually into the map. Inset shows E3-MKLN1-A Kelch β-propeller with Loop-1 and density for Loop-2, extending toward Sub-MKLN1 discoidin domains symmetric with the Sub-MKLN1 Kelch β-propeller binding to E3-MKLN1 shown in Fig. 3b. Loop-2 is indicated by dashed lines. **d** Distal Kelch domain arrangements suggest potential for their interactions between E3-MKLN1-B and Sub-MKLN1-A, and between E3-MKLN1-A and Sub-MKLN1-B. E3-MKLN1-B–Sub-MKLN1-B model derived from fitting domains from E3-MKLN1-A–Sub-MKLN1-A structure (PDB: XXXX) into the density.

**Supplementary Fig. 3.**
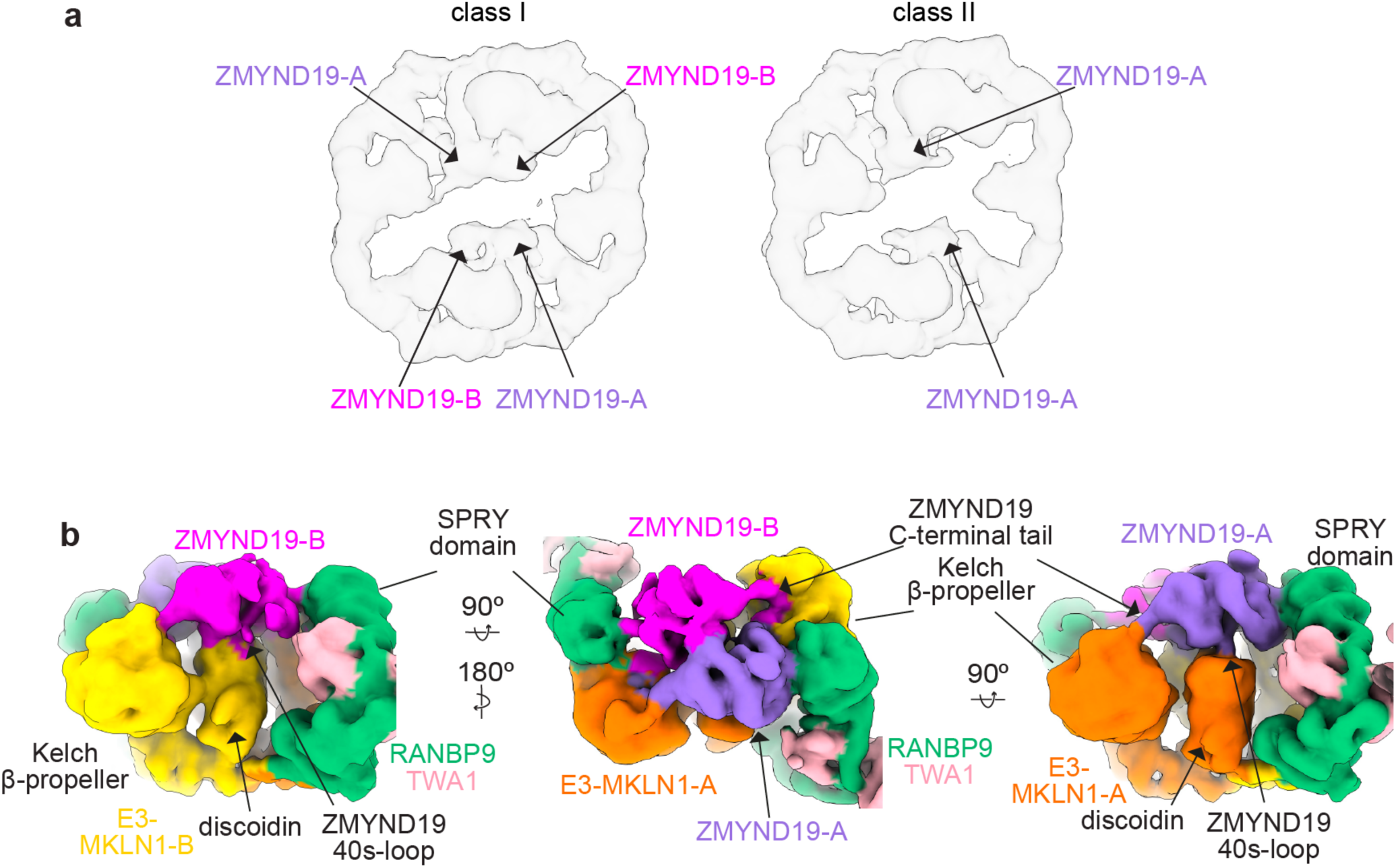
Varying extents of density for ZMYND19 in different 3D classes of complex with CTLH^MKLN1^ and UBE2H∼ubiquitin. **a** Two 3D classes from cryo-EM data processing of CTLH^MKLN1^-UBE2H∼ubiquitin bound to ZMYND19. Density across the entirely of two ZMYND19 molecules (ZMYND19-A and ZMYND19-B) are visible in Class I, whereas the ZMYND domain from ZMYND19-A is only visible in Class II. **b** Three views of cryo-EM map of the MKLN1-RANBP9-TWA1 subcomplex bound to ZMYND19-A and ZMYND19-B, illustrating the asymmetric arrangement of the two ZMYND19 molecules within the complex. Views highlight the ZMYND19 C-terminal tail of both molecules asymmetrically extending toward the Kelch β-propellers from the two E3-MKLN1 protomers, the ZMYND19-A 40s-loop contacting the E3-MKLN1-A discoidin domain, and the ZMYND19 globular ZMYND domain contacting the RANBP9 SPRY domain.

**Supplementary Data Fig. 4.**
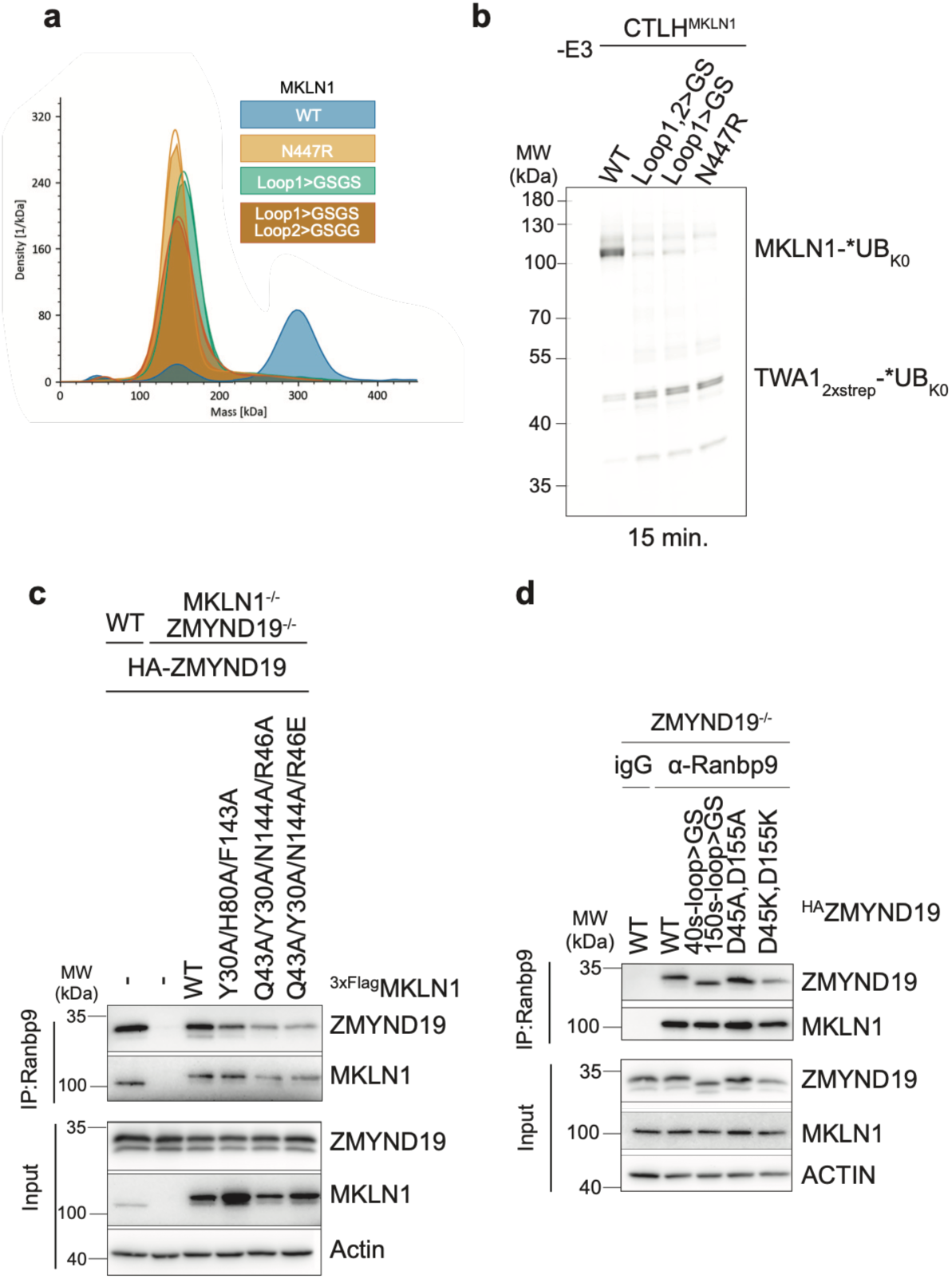
Supporting biochemical and cellular data for CTLH^MKLN1^ substrate recognition. **a** Mass photometry analysis of wild-type MKLN1, N447R, and Kelch domain loop substitution mutants (Loop1>GSGS; Loop1>GSGS and Loop2>GSGG), showing conversion of isolated MKLN1 from a tetramer to a dimer upon mutation. **b** Assays testing effects of MKLN1 Kelch domain loop substitutions (Loop1>GSGS, Loop1,2>GSGS/GSGG, N447R) on MKLN1 ubiquitylation by CTLH^MKLN1^. Increased alternative ubiquitylation of scaffolding subunit TWA1, upon disruption of MKLN1 self-recruitment, is indicated. Assays performed for 15 min. **c** Co-immunoprecipitation via anti-RANBP9 antibody or rabbit IgG as a negative control in MKLN1 and ZMYND19-deficient HEK293 cells co-transfected with HA-ZMYND19 and 3×Flag-MKLN1 WT or discoidin trench mutants, showing retention of ZMYND19 interaction with the CTLH complex. Immunoprecipitates and inputs are shown. Actin serves as loading control. **d** As in **c**, but in ZMYND19 deficient HEK293 cells transfected with WT HA-ZMYND19 or loop substitution mutants (40s-loop>GS and150s-loop>GS; D45A/D155A; D45K/D155K), assessing ZMYND19 loop mutant association with the CTLH complex.

**Supplementary Fig. 5.**
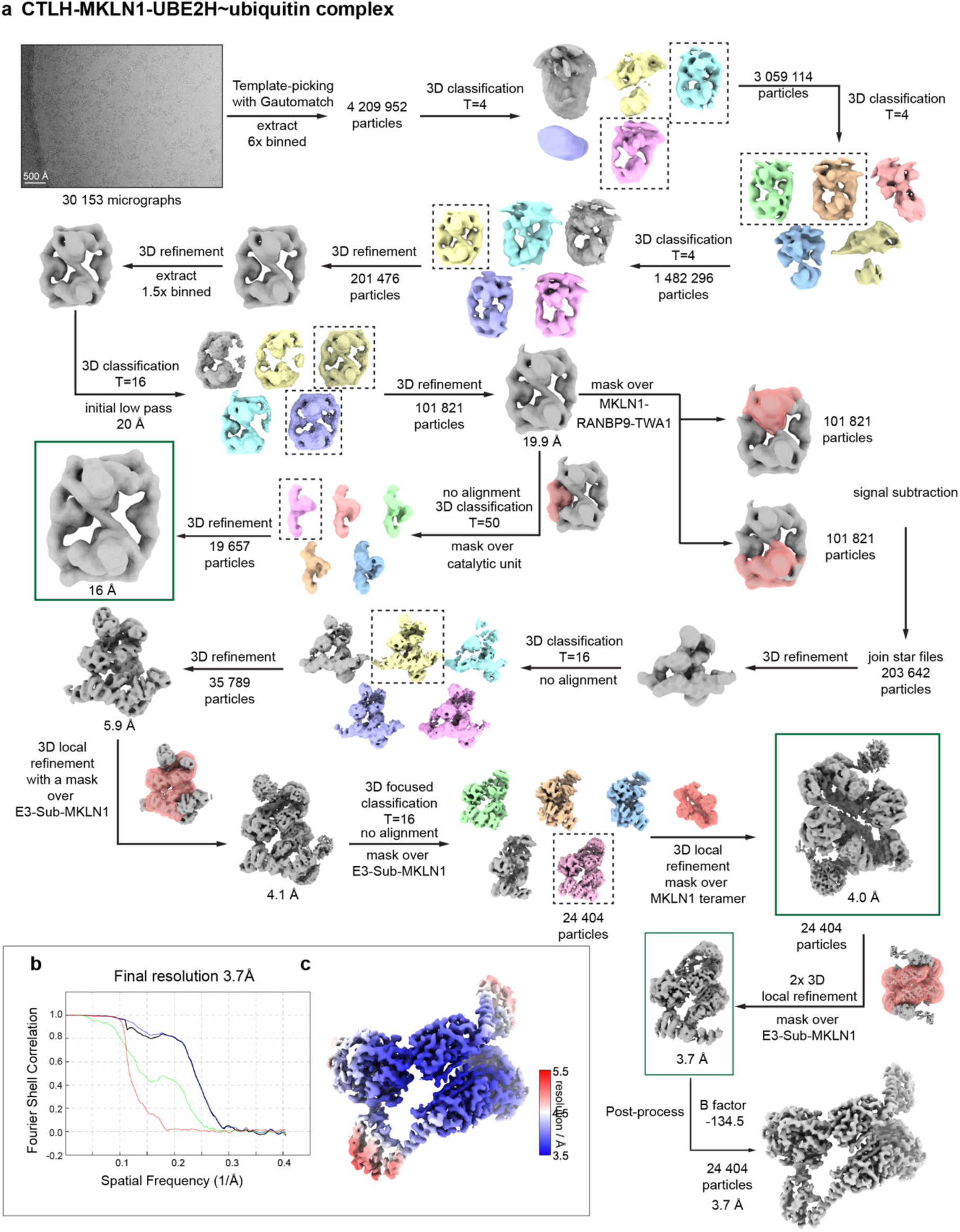
Cryo-EM data processing scheme for the CTLH^MKLN1^ complex. **a** Representative micrograph is shown; scalebar, 500 Å. Data were processed in RELION. Classes selected from classification steps are shown in boxes. Green boxes indicate maps used for Fig. 3b and Supplementary Fig. 2 of this study. Masks are shown on the flowchart arrow when used. **b** Gold-standard Fourier shell correlation at 0.143 cutoff. The final reconstruction reached 3.7 Å resolution. **c** Local resolution estimation.

**Supplementary Fig. 6.**
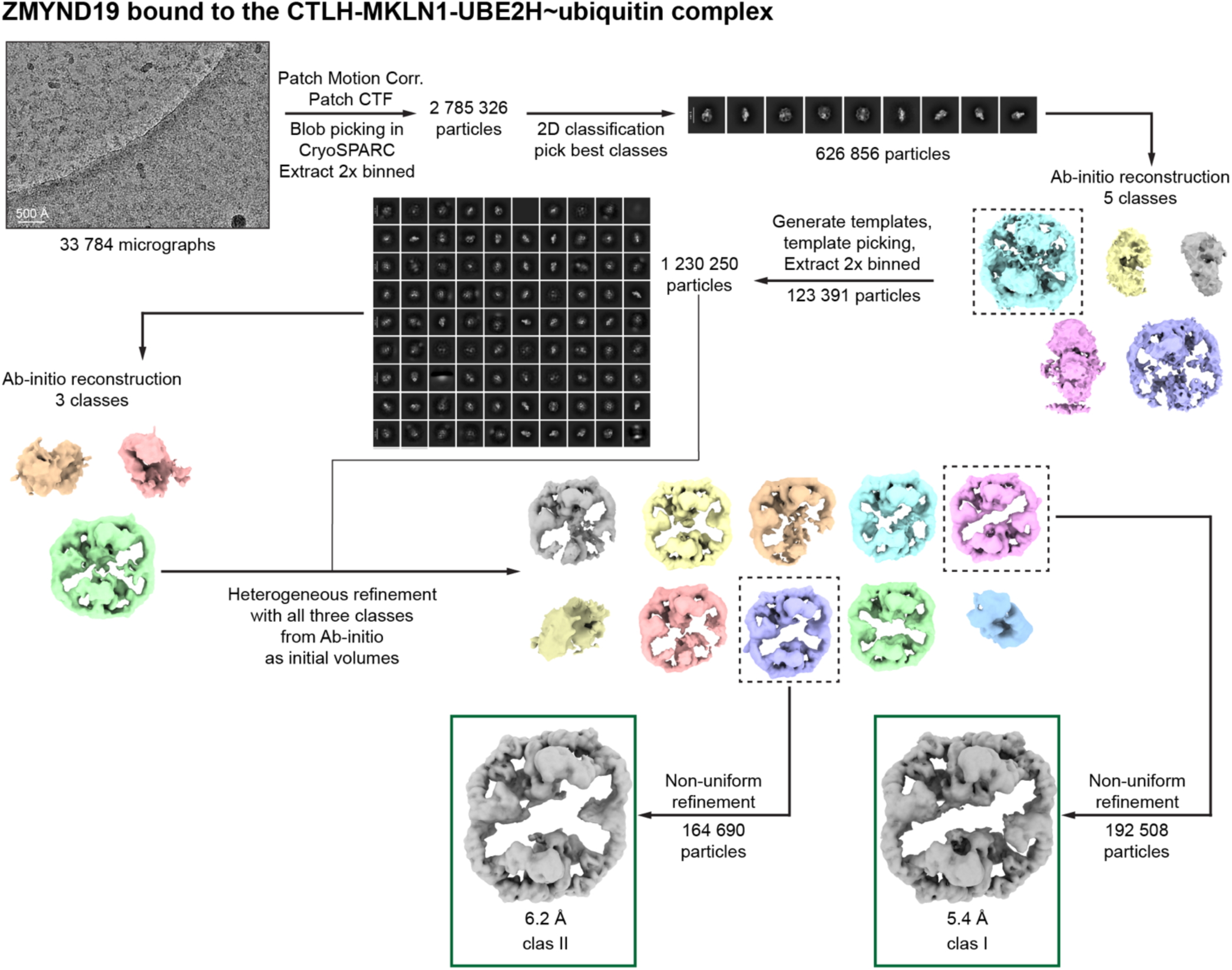
Cryo-EM data processing scheme for the CTLH^MKLN1^-ZMYND19 complex. Representative micrograph is shown; scalebar, 500 Å. Data were processed in cryoSPARC. Templates were generated from Ab-initio model. Green boxes indicate maps used for Fig. 2 (class I) and Supplementary Fig. 3a of this study.

**Supplementary Fig. 7.**
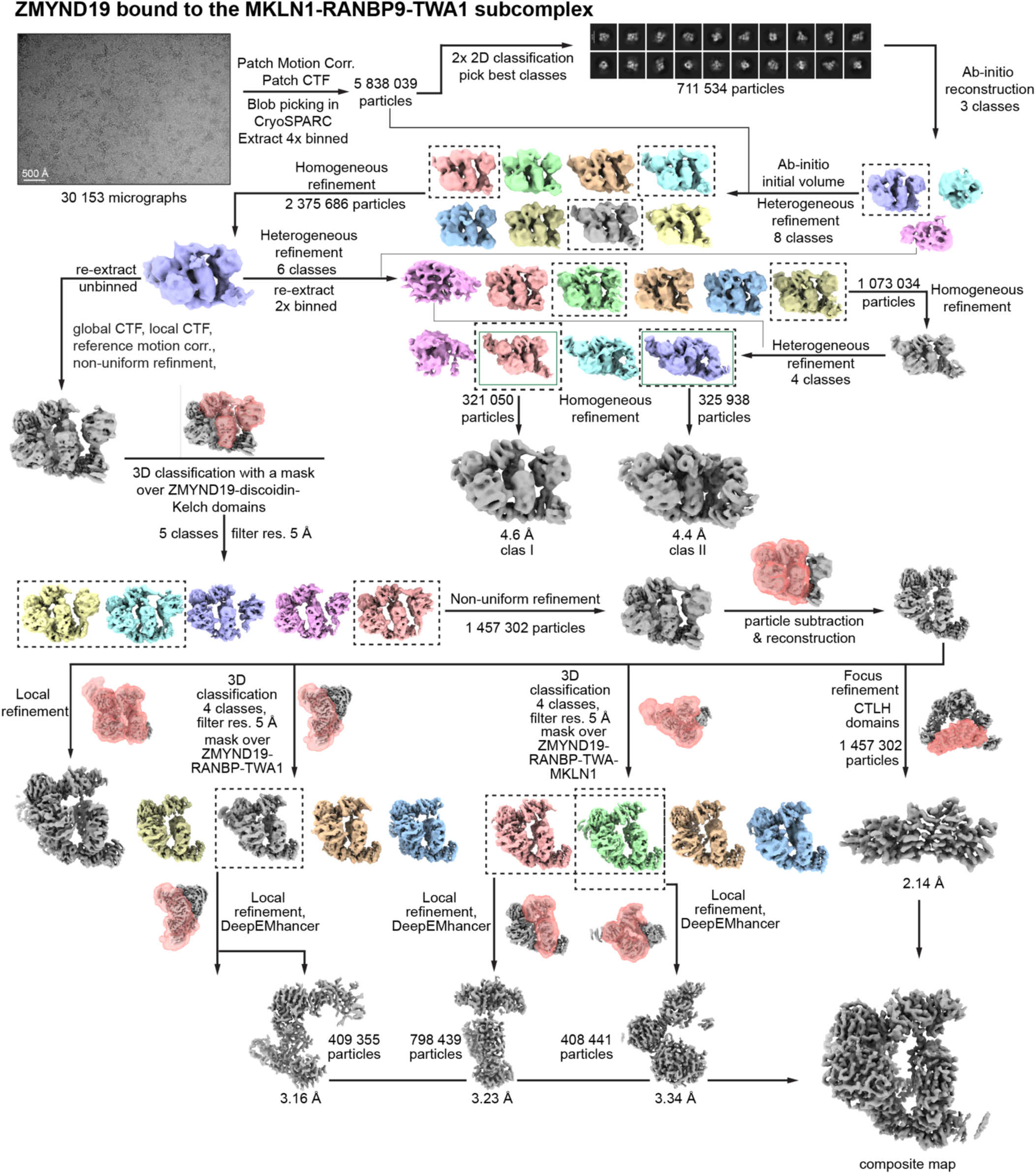
Cryo-EM data processing scheme for the ZMYND19-MKLN1-RANBP9-TWA1 subcomplex. Representative micrograph is shown; scalebar, 500 Å. Data were processed in cryoSPARC. Ab-initio model was directly used for Heterogeneous refinement. Green boxes indicate maps used for Fig. 4a (class I) and Supplementary Fig. 3b of this study. High-resolution maps were achieved through particle subtraction and local refinement with masks. They were used to generate the composite map. Maps used for composite map were sharpened with DeepEMhancer.

**Supplementary Fig. 8.**
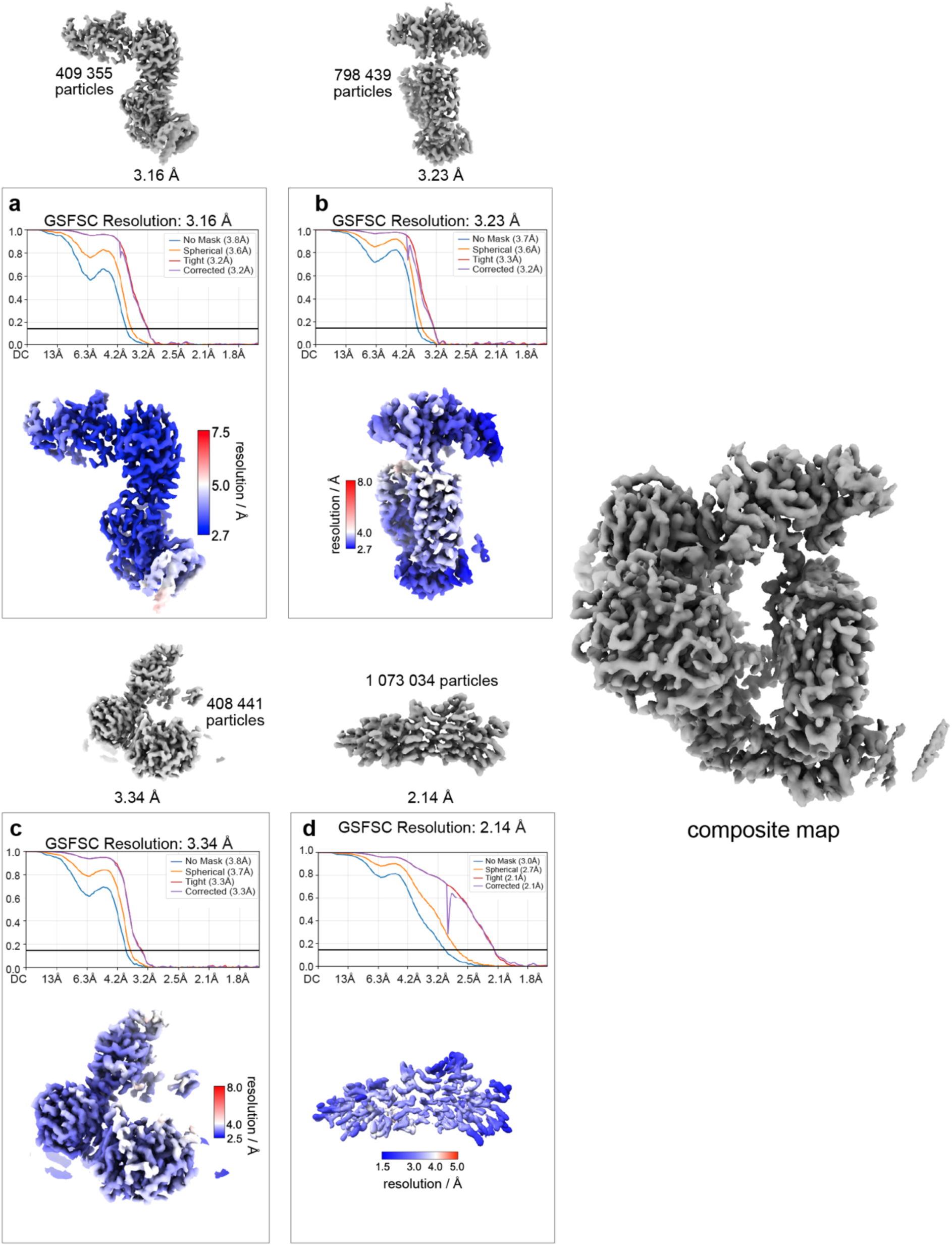
a-d Final reconstructions with particle counts and resolutions displayed on each map used to generate a composite map. Gold-standard Fourier shell correlation (GSFSC) at 0.143 cutoff and local resolution estimation are shown for each sharpened map. The composite map is shown on the right.

**Supplementary Fig. 9.**
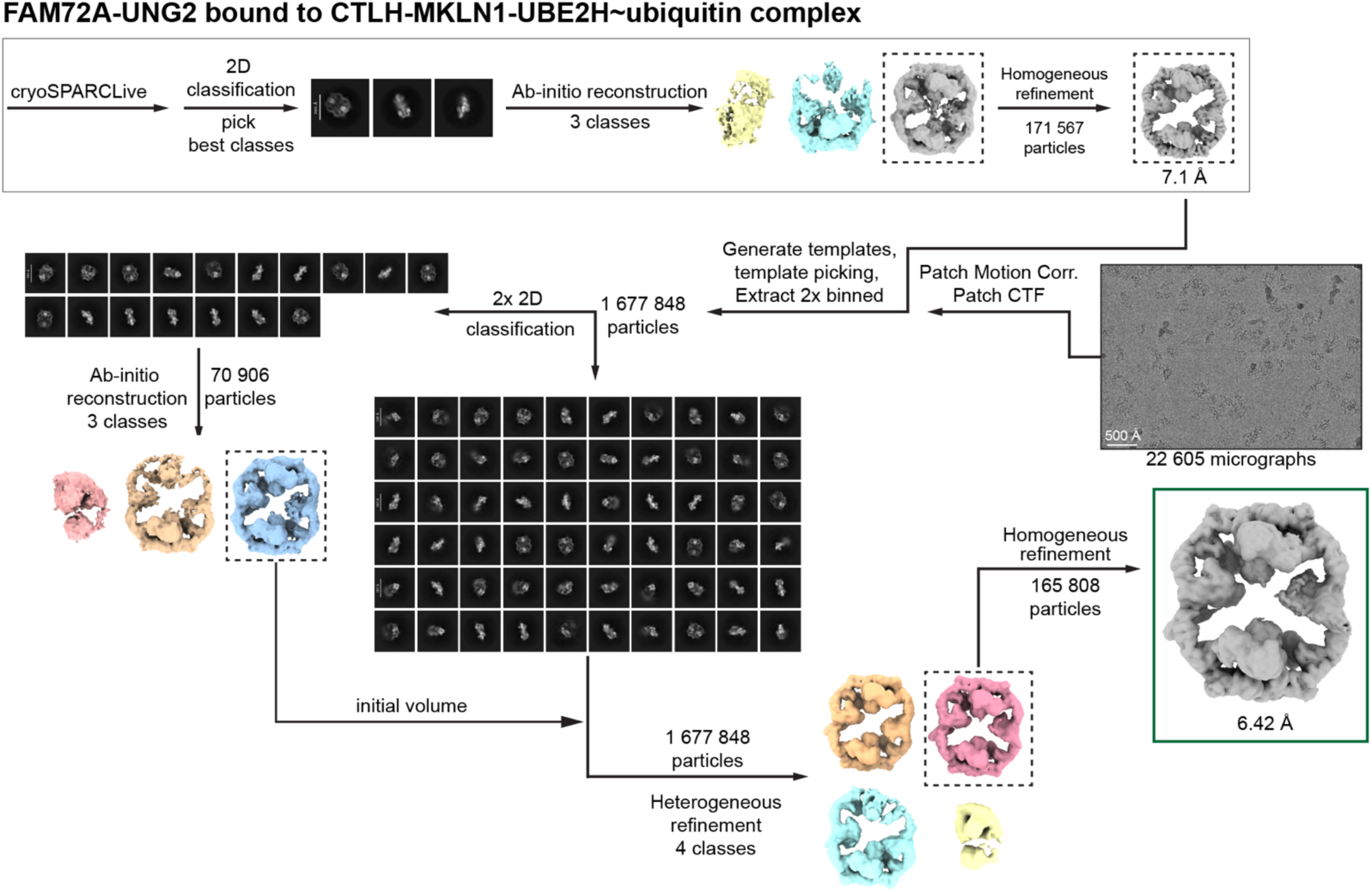
Cryo-EM data processing scheme for the CTLH^MKLN1^-FAM72A UNG2 subcomplex. Representative micrograph is shown; scalebar, 500 Å. Data were processed in cryoSPARC. Templates were generated from cryoSPARC Live processing Ab-initio model. Green box indicates map used for Fig. 2 of this study.

**Supplementary Fig. 10.**
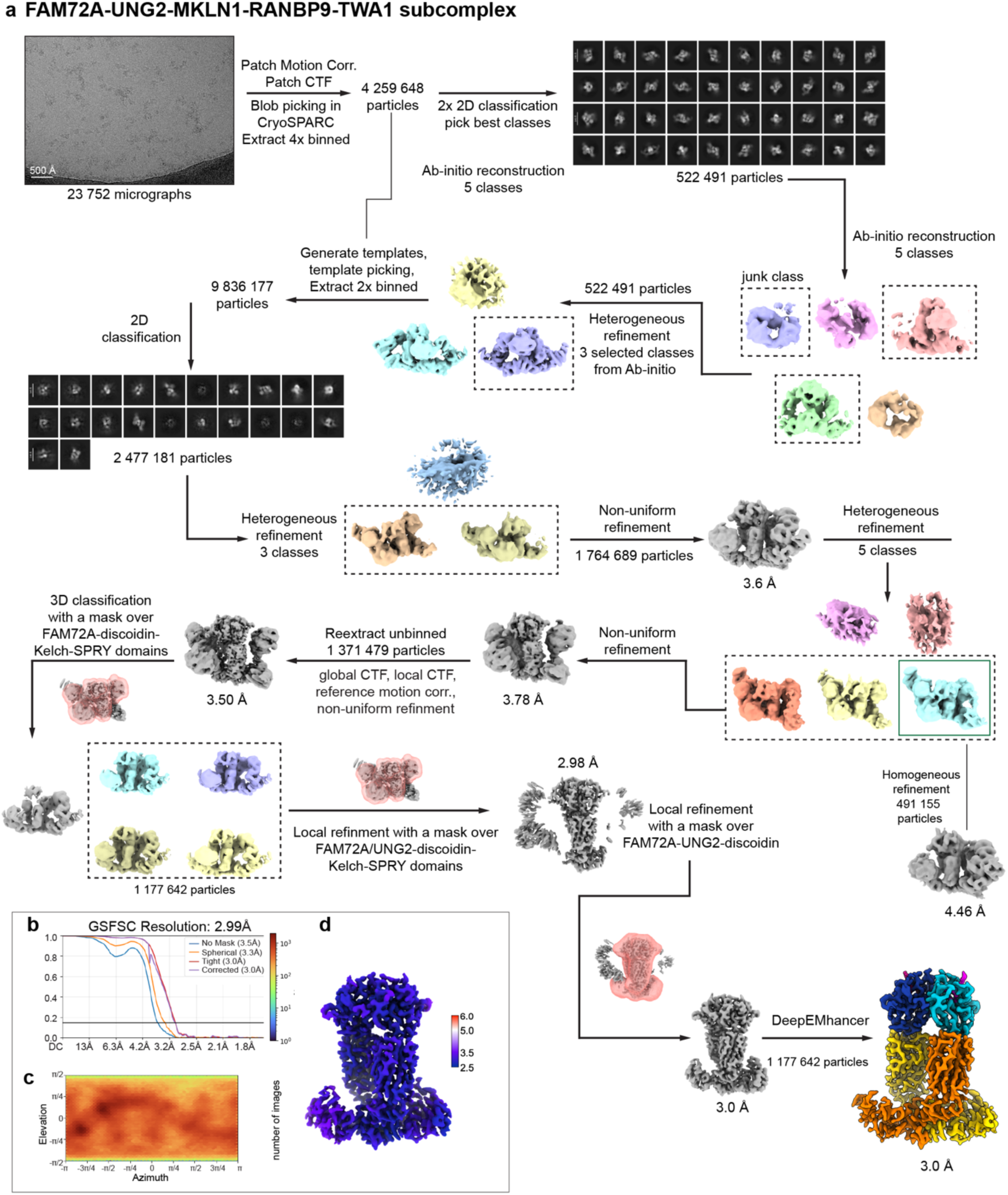
Cryo-EM data processing scheme for the FAM72A-UNG2-MKLN1-RANBP9-TWA1 subcomplex. **a** Representative micrograph is shown; scalebar, 500 Å. Data were processed in cryoSPARC. Classes selected from classification steps are shown in boxes. Heterogeneous refinements contained one junk class. Green box indicates map used for Fig. 5a of this study. **b** The final reconstruction reached 3.0 Å resolution by the gold-standard Fourier shell correlation at 0.143 cutoff and was sharpened with DeepEMhancer. Gold-standard Fourier shell correlation curve. **c** Orientation distribution plot. **d** Local resolution estimation.

**Supplementary Fig. 11.**
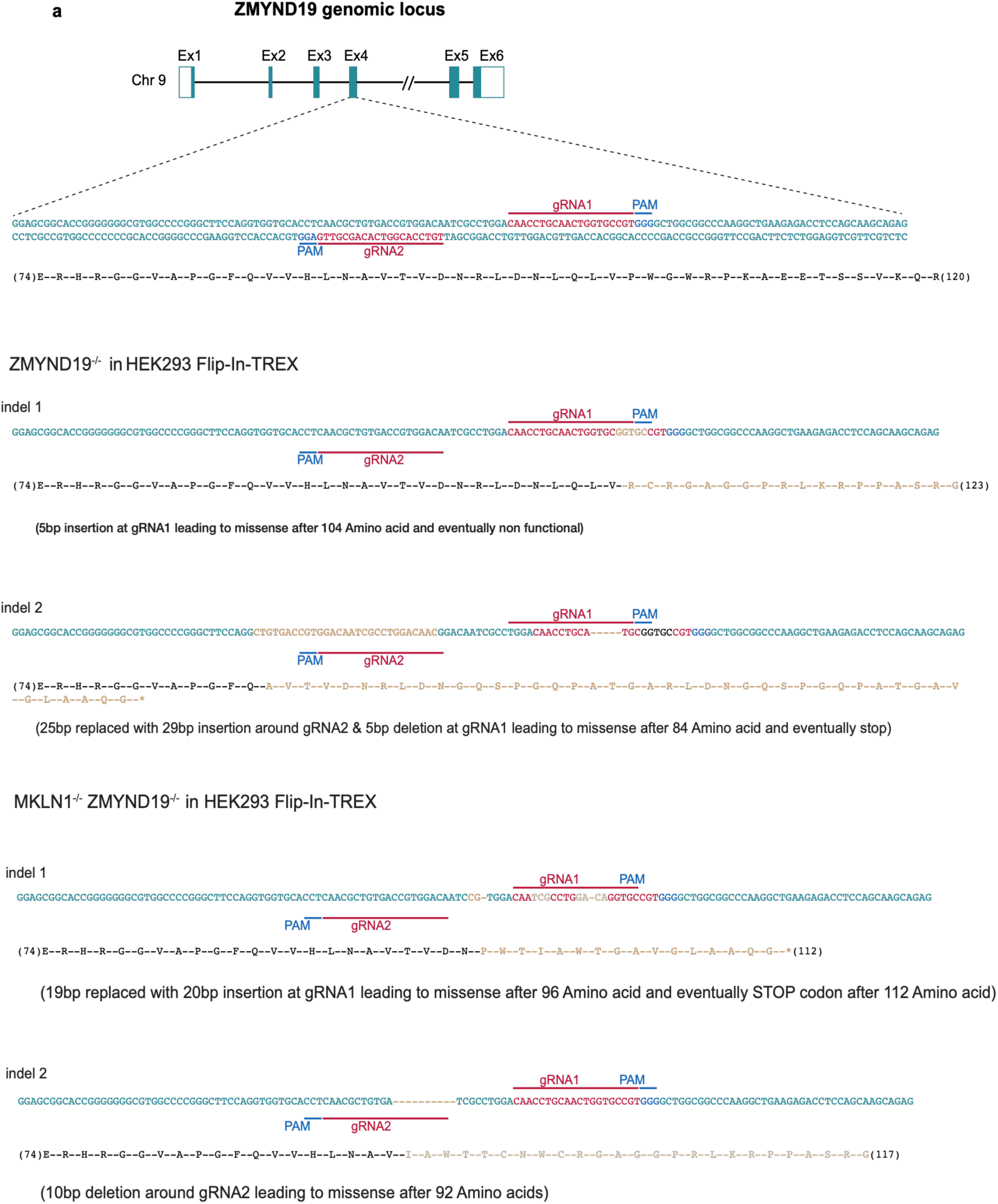
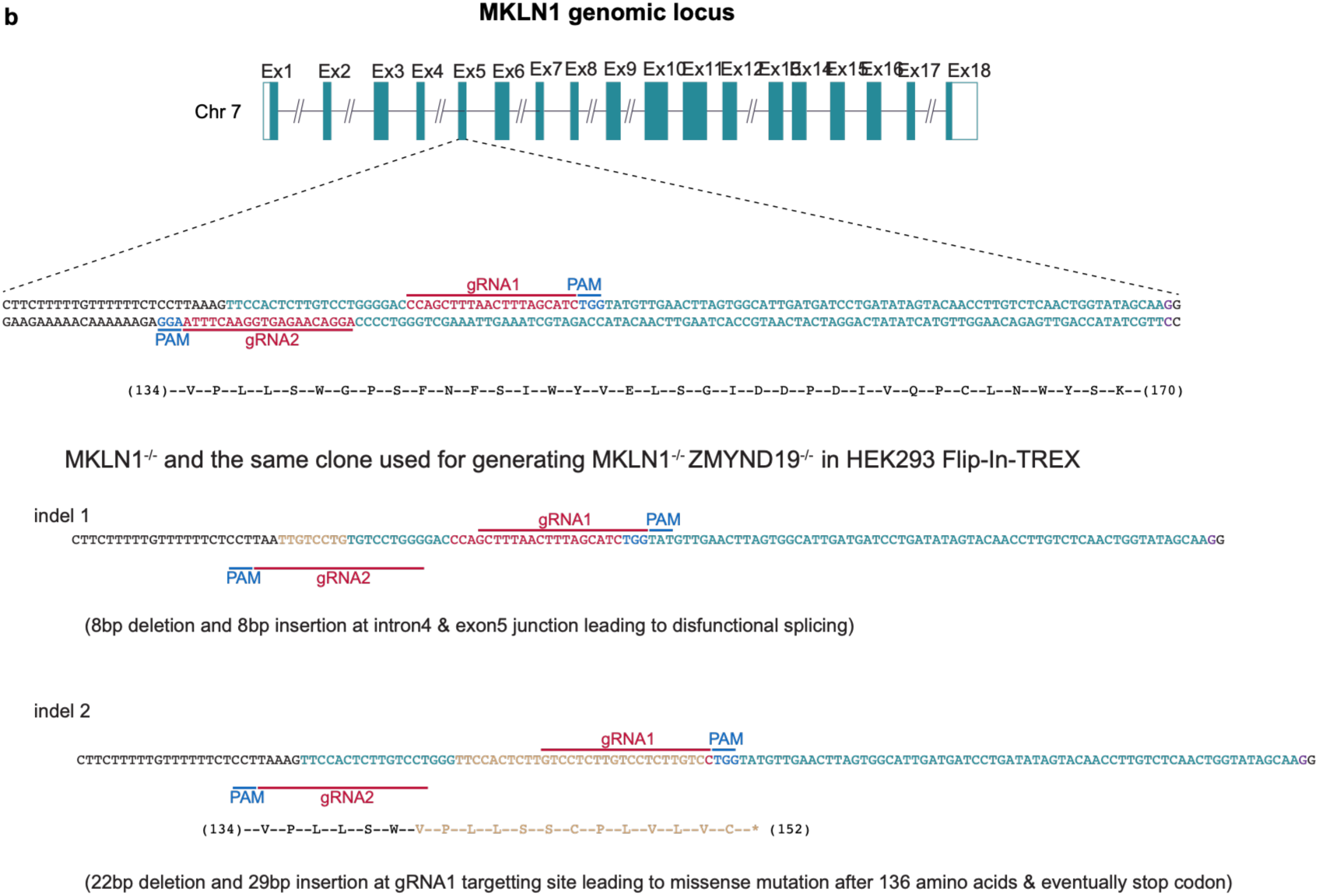
CRISPR Cas9 (nickase) mediated knockout of *ZMYND19*, *MKLN1* and double knockout of *MKLN1* and *ZMYND19*. **a** Schematic representation of genomic locus of *ZMYND19* showing exons structure in genome, location of target guide RNAs (gRNAs) pairs (in red), and PAM sequence (in blue). Insertions/deletions of each knockout clones in Flip-In-TREX HEK293 cells (in orange) and its potential missense effects on ZMYND19 protein expression (in orange). **b** Schematic representation of genomic locus of *MKLN1* showing exons structure in genome, location of target guide RNAs (gRNAs) pairs (in red), and PAM sequence (in blue) analogous to **a**.

